# Cell division in tissues enables macrophage infiltration

**DOI:** 10.1101/2021.04.19.438995

**Authors:** Maria Akhmanova, Attila Gyoergy, Mikhail Vlasov, Fedor Vlasov, Daniel Krueger, Andrei Akopian, Shamsi Emtenani, Aparna Ratheesh, Stefano De Renzis, Daria E. Siekhaus

## Abstract

Migration of cells through diverse tissues is essential for development, immune response and cancer metastasis ^1–3^. To reach their destination, cells must overcome the resistance imposed by complex microenvironments, composed of neighboring cells and extracellular matrix (ECM)^4–6^. While migration through pores and tracks in ECM has been well studied ^4,5,7^, little is known about cellular traversal into confining cell-dense tissues. Here by combining quantitative live imaging with genetic and optogenetic perturbations we identify a crucial role for cell division during cell migration into tissues. We find that normal embryonic invasion by *Drosophila* macrophages between the ectoderm and mesoderm^8,9^ absolutely requires division of an epithelial ectodermal cell at the site of entry. Dividing ectodermal cells disassemble ECM attachment formed by Integrin-mediated focal adhesions next to mesodermal cells, allowing macrophages to move their nuclei ahead and invade. Decreasing or increasing the frequency of ectodermal division correspondingly either hinders or promotes macrophage invasion. Reducing the levels of focal adhesion components in the ectoderm allows macrophage entry even in the absence of division. Our study demonstrates the critical importance of division at the entry site to enable *in vivo* cell invasion by relieving the steric impediment caused by focal adhesions. We thus provide a new perspective on the regulation of cellular movement into tissues.

## Introduction

Cell dissemination into tissues is fundamentally important for the formation and maintenance of complex organisms. During vertebrate development, neural crest cells move into tissues to form bone, neural, glial, endocrine and connective tissue^3,10^ and immune cells enter organs to establish residence and regulate tissue function^11–13^. During inflammatory responses immune cells infiltrate tissues to attack or engulf pathogens^14^. Finally, during the metastatic spread that underlies mortality, cancer cells traverse into other organs^2,15^. These various embryonic and adult environments contain closely packed cells adherent to each other or to the ECM that lies between them^16–18^. Despite its crucial importance for development, immunity, and cancer, invasion into such cell-dense tissues is poorly understood^4,6,7,15^.

Macrophages invade tissues from early on in development to establish residency, enabling immune scavenging of molecules, debris, dying cells or invading pathogens^12,19^. In the early *Drosophila* embryo macrophages follow guidance cues^20^ and invade at a particular location into the germband between the ectoderm and the mesoderm, which at this stage are juxtaposed and separated only by a thin layer of extracellular matrix (ECM)^8,9^(Fig. 1a, b). The edge of these tissues represents a barrier that macrophages must overcome in order to invade, and the first macrophage requires around 20 min to enter after reaching this location. Macrophage-specific programs, not involving proteolytic ECM degradation^8^, are known to affect the efficiency of macrophage entry^21–23^. However, how the dynamics and properties of surrounding cells influence macrophage tissue invasion remains unclear^9^. We took advantage of the fact that during early *Drosophila* development macrophage invasion occurs at a precise time and place to approach this question in an *in vivo* context.

**Fig. 1.**
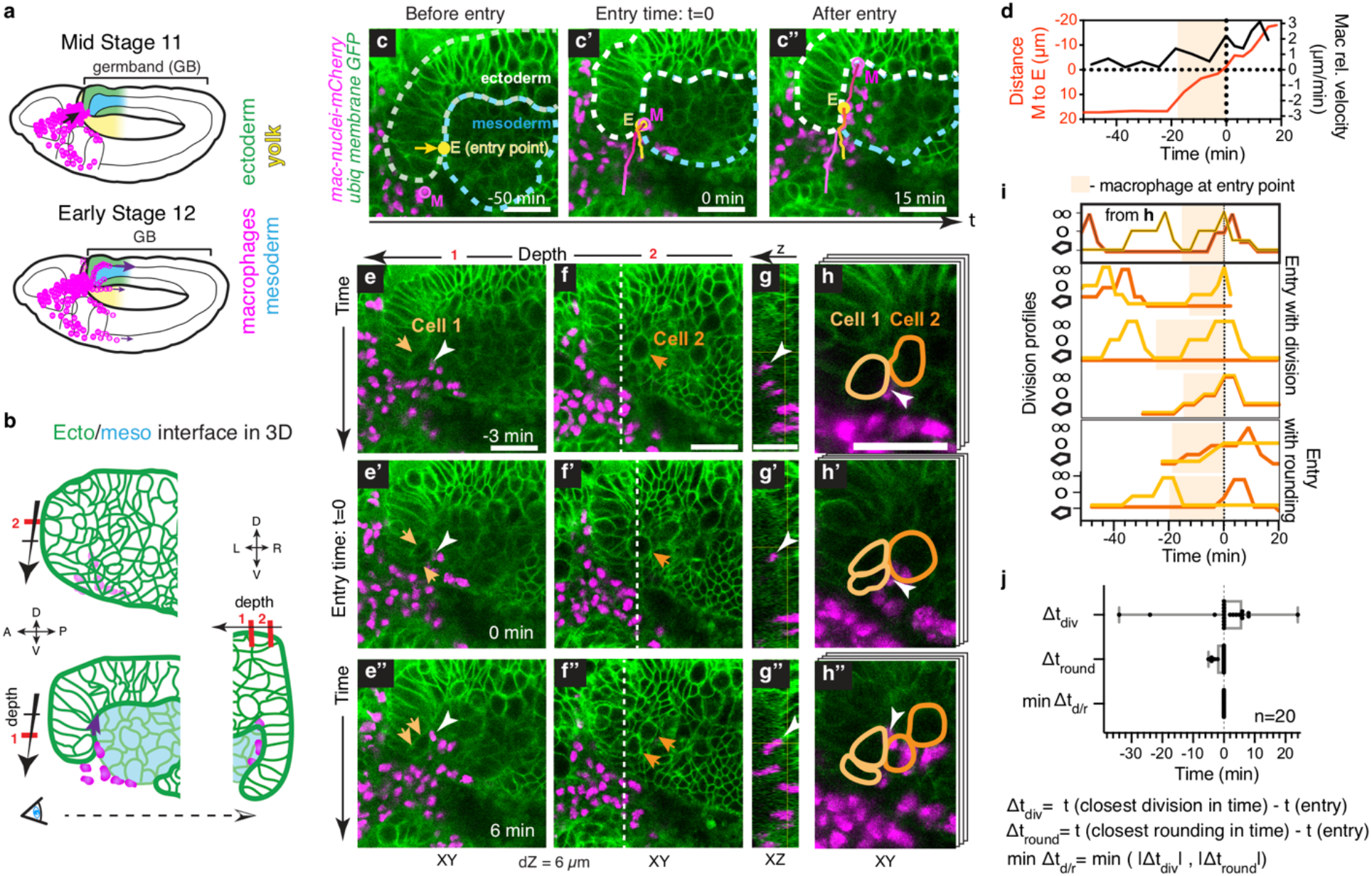
Macrophage invasion correlates with ectodermal cell division at the entry point. **a,** Lateral embryo schematic showing macrophages entering the germband (GB) and migrating along the ectoderm and mesoderm interface. **b**, Lateral germband zoomed cross-sections: (1) deeper, (2) closer to the surface. Orthogonal view at right. **c,** Tracks trace the location over time of the first entering macrophage (M, magenta) and the macrophage entry point (E, yellow). **d,** Graph of the distance (red) between the first macrophage’s nucleus (M) and the tissue entry point (E) and the corresponding velocity of M relative to E (black). **e,** Stills showing an entering macrophage (white arrowhead) and the dividing adjacent Cell 1 (arrows) at the entry point. Z-slice position (1) in (**b**). **f,** Stills 6 μm higher than (**e**), showing ectodermal Cell 2 (arrows) undergoing mitotic rounding and division above the entering macrophage. Z-slice position (2) in (**b**). **g,** Orthogonal view at dashed white line in (**f**). **h,** Maximum intensity projection of zoomed slices in (**e, f**) with Cell 1 and 2 are outlined. **i,** Division profiles in time of Cell 1 and Cell 2 outlined in (**h**) and similar profiles from 5 other embryos. Bottom, middle and top positions on the y-axis indicate that a cell is in interphase, undergoing mitotic rounding, or dividing (cell shapes on y-axis). Macrophages enter germband at time=0. **j,** Quantification of the time difference between macrophage entry and the division (top), or rounding (middle) of an adjacent cell at the entry point and the minimum of these two values (bottom). Box plots: 25^th^-75^th^ percentiles, whiskers: max-min values. Scale bars: 20μm. (**c-h**) genotype: *Resille::GFP*, *DE-Cad::GFP*; *srpHemo-H2A::3xmCherry*.

## Results

### Macrophages invade into cell-packed tissue at a defined location

To visualize tissue invasion we imaged tissues labelled with a plasma membrane marker and tracked the nuclei of migrating macrophages (Fig. 1c, Video 1). The nucleus is the stiffest organelle and is used by cells as a ruler for pathfinding in narrow spaces^7^. The macrophage entry point lies in the acute angle formed by the basal side of the ectoderm and the mesoderm surface (Fig. 1b,c). To assess entry quantitatively, we plotted the distance between the first entering macrophage and the entry point in the dorso-ventral (vertical) direction over time. We defined entry as occurring when the first macrophage crossed the horizontal plane drawn through the entry point causing the separation of the previously adjacent ectodermal and mesodermal cells. Analysis of macrophage velocity relative to the entry point shows that the first macrophage displayed a low-speed phase, i,e. a pause, as it first contacted the surrounding cells at the entry point, and thereafter a high-speed phase, i.e. a jump (Fig. 1d, Fig. S1a-d). Subsequently the macrophage nucleus is seen between the ectodermal and mesodermal cells. We used the time of the jump to precisely define the moment of entry.

### Macrophage entry occurs next to a dividing ectodermal cell

We noticed that as the first macrophage entered, one or two ectodermal cells adjacent to the entry point had become round, a shape change that starts at the beginning of the mitotic process of cell division^24^, or had progressed to become two connected smaller rounded spheres, showing the cell was in the final phase of division (Fig. 1e-h). During entry the first macrophage touched two ectodermal cells, as shown with outlines in Fig. 1h. Corresponding plots of the mitotic rounding and division of these cells over time are shown in Fig. 1i (upper graph). By aligning the division profiles of 20 movies using the timepoint when the first macrophage moved between the two tissues, we observed that macrophage entry always correlated with the division (Fig. 1i, Fig. S1e,g) or rounding (Fig. 1i, Fig. S1f,h) of a flanking ectodermal cell. We calculated the time difference between macrophage entry and the most recent mitotic rounding or division. This number was always 0 (Fig. 1j), showing a complete concordance between macrophage entry and one of these two mitotic events. We quantified how much time the ectodermal cells at the entry point spent in each of the three following categories: 1) at least one of the two cells is dividing, 2) at least one cell is rounding up 3) none of these cells are mitotic, the interdivision phase (Fig. S1i). If macrophage invasion occurred randomly, one would expect to find macrophages entering during the interdivision phase in more than half of the embryos; instead, we never observed entry at this time (Fig. S1j).

We found a further indication of the importance of ectodermal division by examining the two bilaterally symmetrical entry points on the sides of the hindgut that lie within a dorsally viewed embryo. Macrophages arrived at these two locations almost simultaneously, however they did not enter at the same time if ectodermal divisions at both points did not occur synchronously (Fig. S2). Finally, we also examined if division of the adjacent mesodermal cells correlated with macrophage entry and found that it did not (Fig. S3). We therefore conclude that in wild type embryos macrophages enter into the germband tissue only when an adjacent ectodermal cell is dividing, suggesting that this is a permissive event.

### Inhibition of cell division blocks entry

Next, we examined if ectodermal division is required for macrophage entry. We injected embryos with the drug dinaciclib, which inhibits Cdk1 and other CD-kinases, and effectively stops the progression of cell division (Fig. 2a, Fig. S4a). The density of rounded cells in the ectoderm was reduced by 83% after dinaciclib injection (Fig. 2b). Macrophages remained motile and moved directionally (Fig. S4b-c), however, they almost completely failed to enter into the germband (Fig. 2c-e’). We quantified the number of macrophages inside the germband at the time of germband retraction, which is ~60 min after macrophages arrived at the entry point. In control embryos, ~40 macrophages had moved inside the germband by this time (Fig. 2c, c’, e). However, in the great majority of embryos treated with dinaciclib, no macrophages (50% of embryos) or only a few (up to 10 in 30% of embryos) entered the germband (Fig. 2d-e). Live imaging revealed that in embryos in which a few macrophages entered into the tissue, they moved in at the usual entry location next to a round ectodermal cell that did not progress to complete division for a long period of time (Fig. 2f, Video 2, Fig. S4d-g). Thus, pharmacological inhibition of cell division resulted in macrophages not invading at all or entering next to the remaining mitotically rounded ectodermal cells (Fig. 2g).

**Fig. 2.**
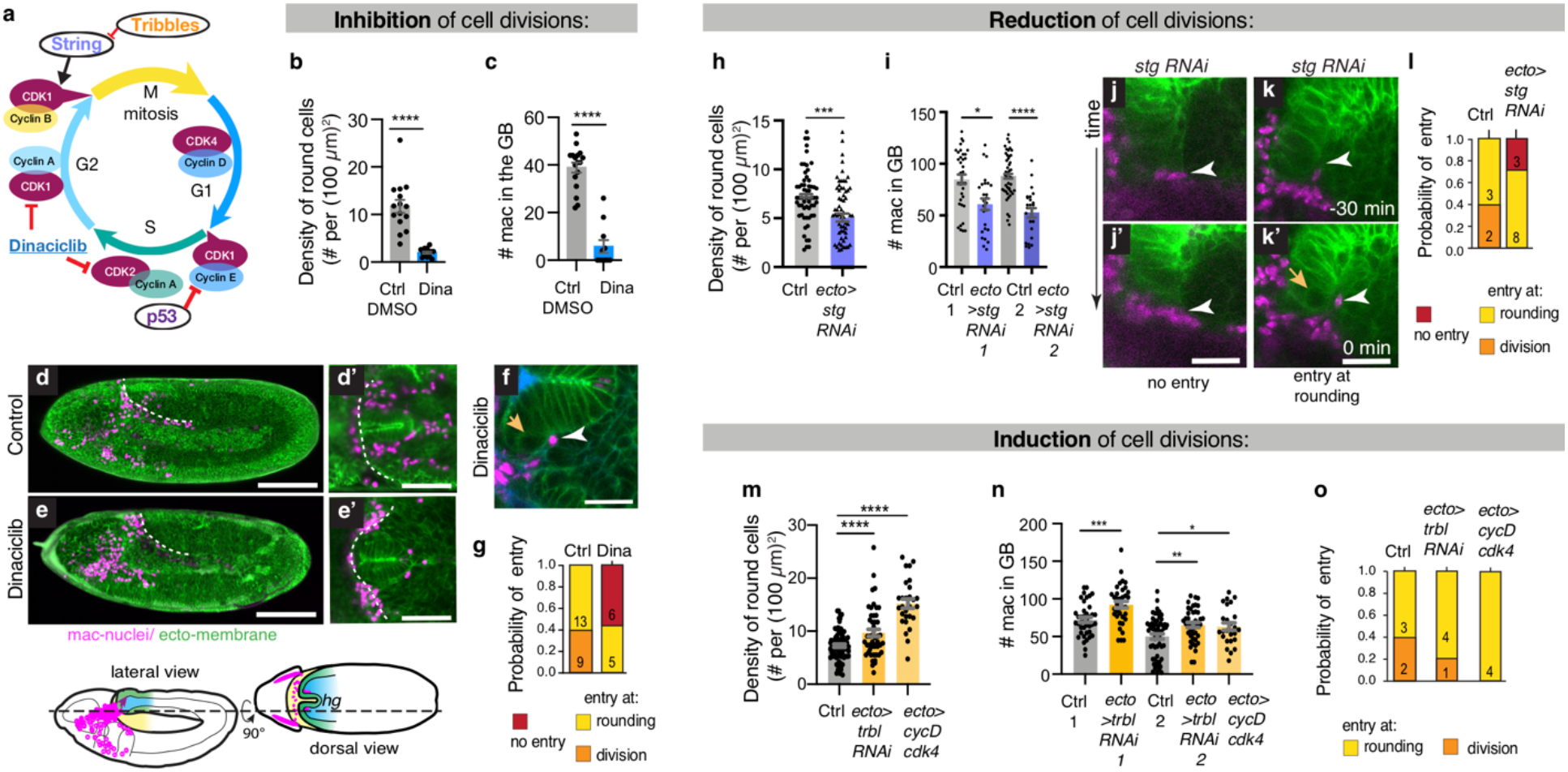
Macrophage entry requires ectodermal mitotic rounding. **a,** Cell cycle regulation schematic. Dinaciclib inhibits CDKs. **b,c,** Rounded cell density in the germband tissue at macrophage entry time (**b**) and macrophage numbers inside the germband as germband retraction starts (**c**) in control DMSO-injected embryos (n=15, 17) and dinaciclib-injected embryos (n=14, 11). N=4 experiments. ****p<0.0001, Mann-Whitney test. **d-e,** Maximum intensity projections of stage 12 embryos injected 60 min earlier with (**d-d’**) DMSO as a control or (**e-e’**) dinaciclib. (**d, e**) lateral, (**d’, e’**) dorsal view. Schematics below. **f,** Time-lapse zoomed image of the first macrophage (white arrowhead) entering the germband in a dinaciclib-injected embryo next to a rounded ectodermal cell at the entry site (orange arrow). **g,** Probability distribution of macrophage entry. **h,** Rounded cell density in the germband tissue in control embryos (N=7 embryos, z=5-10 planes per embryo, n=58) and those expressing *stg RNAi* in the ectoderm (N=8 embryos, z=5-10 planes per embryo, n=70); means ±s.e.m, ***p=0.0001, unpaired t-test. **i,** Macrophage numbers inside the germband; means± s.e.m, *p=0.046, ****p<0.0001 one-way ANOVA with Tukey, n=48, 26, 46, 25. **j-k,** Timelapse images of *ecto*>*stg RNAi* embryos. (**j**) No macrophages enter and no entry point division or rounding occurs for >60 min. (**k**) Macrophages enter next to rounded cell. **l,** Probability distribution of macrophage entry. **m**, Rounded cell density in the germband tissue in control embryos (n=7 embryos, z=5-7 planes per embryo) and those expressing either *trbl RNAi2* (n=8) or *cycD* and *cdk4* in the ectoderm (n=5, z=5-7 planes per embryo); means± s.e.m, ****p<0.001, one-way ANOVA with Tukey. **n,** Macrophage numbers inside the germband, means± s.e.m; control 1 (n=40), *ecto*>*trbl RNAi1* (n=36), ***p=0.0007 t-test; control 2 (n=60), *ecto*>*trbl RNAi2* (n=43), *ecto*>*cycD,Cdk4* (n=25), **p<0.0017 *p=0.0344, one-way ANOVA with Tukey. **o,** Probability distribution of macrophage entry. Scale bars: 100μm (**d**, **e**); 50μm (**d’**, **e’**); 20μm (**f**, **j-k**).

To confirm these results, we inhibited the division frequency only locally, expressing RNAis against a positive regulator of mitosis, cdc25 (*string*, *stg*) in the ectoderm (Fig. 2a) which reduced the density of rounded cells in this tissue before macrophage entry by ~30% (Fig. 2h). This corresponds to a lower division frequency also at the entry point. The number of macrophages that could penetrate into the germband was ~25-40% lower upon the ectodermal expression of two different *stg RNAi*s compared to the control (Fig. 2i, Fig. S4h). Live imaging identified embryos expressing these RNAis in which no divisions occurred in the germband edge for a long time, and no macrophages entered (Fig. 2j, j’, Video 3 left). When we did detect entry in other embryos, it was always adjacent to a mitotic ectodermal cell at the normal location (Fig. 2k, k’, Video 3 right). Overexpression of the negative cell cycle regulator *p53* in the ectoderm decreases division density by 30% (Fig. 2a, Fig. S4i) and also led to a 35% decrease in macrophage numbers in the germband (Fig. S4j). We conclude that reducing the frequency of ectodermal cell division reduces macrophage entry, while maintaining the entry-mitotic rounding correlation (Fig. 2l, Fig. S4k).

### Induction of cell division prompts entry

Having established that cell division is required for macrophage invasion, we tested whether increasing the rate of mitosis in the ectoderm would cause the opposite result. To this end, we analyzed embryos expressing RNAi against a negative regulator of mitosis, *tribbles*, and those overexpressing the G1 progression regulators *CyclinD* and *Cdk4* (Fig. 2a) in the ectoderm. These treatments increased the density of rounded cells before macrophage entry by 25-80% (Fig. 2m), and led to an overall increase in macrophage numbers in the germband by ~20% compared to the control (Fig. 2n, Fig. S4l). Macrophages always entered at their usual position and next to a mitotic ectodermal cell (Fig. 2o, Fig. S4m, Video 4). These results argue that the timing of ectodermal divisions is normally the rate-limiting factor for macrophage invasion.

### Mitosis disassembles focal adhesions

We next asked how ectodermal mitotic rounding facilitates macrophage entry. As the cell cortex stiffens two-fold during mitotic rounding^24^, we tested if this might help macrophages to move ahead. We increased cortex contractility through optogenetic recruitment of the Rho1 exchange factor RhoGEF2 to the plasma membrane (Fig. S5a)^25^, inducing rounding of ectodermal cells in a small region (50×40×20 μm) of the ectodermal edge including cells at the entry site (Fig. S5b-d). However, this caused no change in the timing of macrophage invasion (Fig. S5e-g). These results suggest that an increase in cortical tension of ectodermal cells alone is not sufficient to enhance macrophage entry and that other mechanisms explain the link between ectodermal division and macrophage invasion.

Focal adhesions have been observed to disassemble during vertebrate mitotic rounding *in vitro*^26,27^. We examined their *in vivo* temporal and spatial dynamics in the germband to determine if they could influence macrophage entry. The ectoderm faces the mesoderm with its basal side, forming focal adhesions that bind through Integrin to the underlying thin layer of Laminin between the two tissues (Fig. 3a,b)^9^. We first visualized these focal adhesions in fixed embryos with antibody staining against Integrin, which localized to dot-like adhesive structures at the ectodermal-mesodermal interface (Fig. S6a). Live imaging using fluorescently tagged intracellular focal adhesion components Vinculin or Talin^28^, also revealed a dotted pattern along this interface (Fig. 3b,c, Fig. S6b). Once an ectodermal cell started to round, these focal adhesions visualized by Vinculin-mCherry gradually disappeared, leaving only one peak remaining in the middle of the basal side (Fig. 3d,e, Fig. S6c-g). Thereafter this last focal adhesion peak gradually flattened (Fig. 3e,f).

**Fig. 3.**
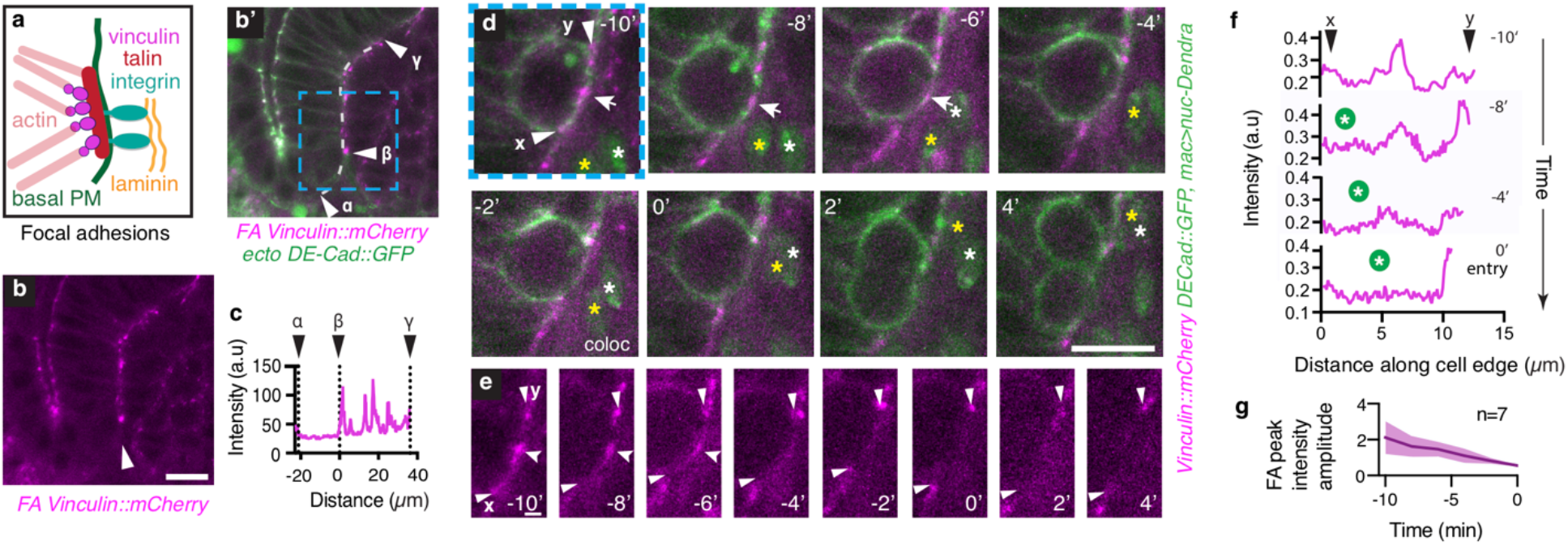
Macrophages enter when focal adhesions (FA) disassemble during ectodermal mitotic rounding. **a,** Scheme of molecular composition of ectodermal focal adhesions (FA). **b,** FA in ecto-meso interface visualized by Vinculin::mCherry. **b’,** Merge of (**b**) with image of ectodermal cell membranes labeled by DE-Cad::GFP. White line indicates basal side of ectoderm abutting mesoderm. **c,** Vinculin::mCherry intensity along ecto-meso interface from (**b,b’**). Point *β*: future macrophage entry point. **d,** Timelapse imaging of ectodermal FAs during mitotic rounding and subsequent macrophage entry. Macrophage nucleus (yellow and white stars) labeled with *srpHemo-H2B::Dendra*. **e,** Vinculin::mCherry on the basal side of mitotic cell, cutout from (**d**). **f,** Vinculin::mCherry intensity along the basal side of rounding cell in (**d,e**) from point **x** to **y** over time. Green circle: first macrophage nucleus. **g,** Amplitude of FA peak over time. Mean ± s.d (shading). Scale bars: 10μm (**b**-**b’, d**), 2μm (**e**).

Focal adhesion disassembly did not depend on the presence of macrophages and happened in every basally dividing cell in the ectoderm (Fig. S6e-h, Video 5). Importantly, imaging ectodermal mitotic cells at the entry point revealed that macrophage entry always occurred after the last adhesion spot disassembled (Fig. 3e, Video 6, Fig. S6i,j). In cases in which entry occurred next to an ectodermal cell that had just divided, mitosis was completed faster than macrophage nuclear translocation, however new focal adhesions were not established before macrophage entry. We conclude that focal adhesion disassembly at the entry point is required for the macrophage nucleus to penetrate between the ectodermal and mesodermal cells.

### Focal adhesion reduction controls entry

Finally, we tested if ectodermal adhesion is the main factor hindering macrophage invasion between the ectoderm and mesoderm. We reduced focal adhesion components in the ectoderm by *RNAi* (Fig. S7) and found that the number of macrophages entering the germband was higher than in the control (Fig. 4a-d, Fig. S7a,b). Importantly, through live imaging we directly observed that the first macrophage could enter without ectodermal rounding or division in embryos with lower levels of *β*-Integrin or Vinculin in the ectoderm (Fig. 4e,f, Video 7). Cumulative probability graphs based on these movies show that macrophage entry occurred frequently without ectodermal division upon ectodermal knockdown of focal adhesion components in contrast to the wild type control in which this was never observed (Fig. 4f). Even in embryos in which dinaciclib injection blocked cell cycle progression, expression of *RNAi* against *β*-Integrin in the ectoderm resulted in macrophages entering in the absence of rounding or dividing cells at the entry point (Fig. S7k). We therefore conclude that normally ectodermal focal adhesions inhibit macrophages from entering into the germband tissue and that ectodermal cell division opens the door for macrophages to move in by disassembling these adhesions during mitotic rounding.

**Fig. 4.**
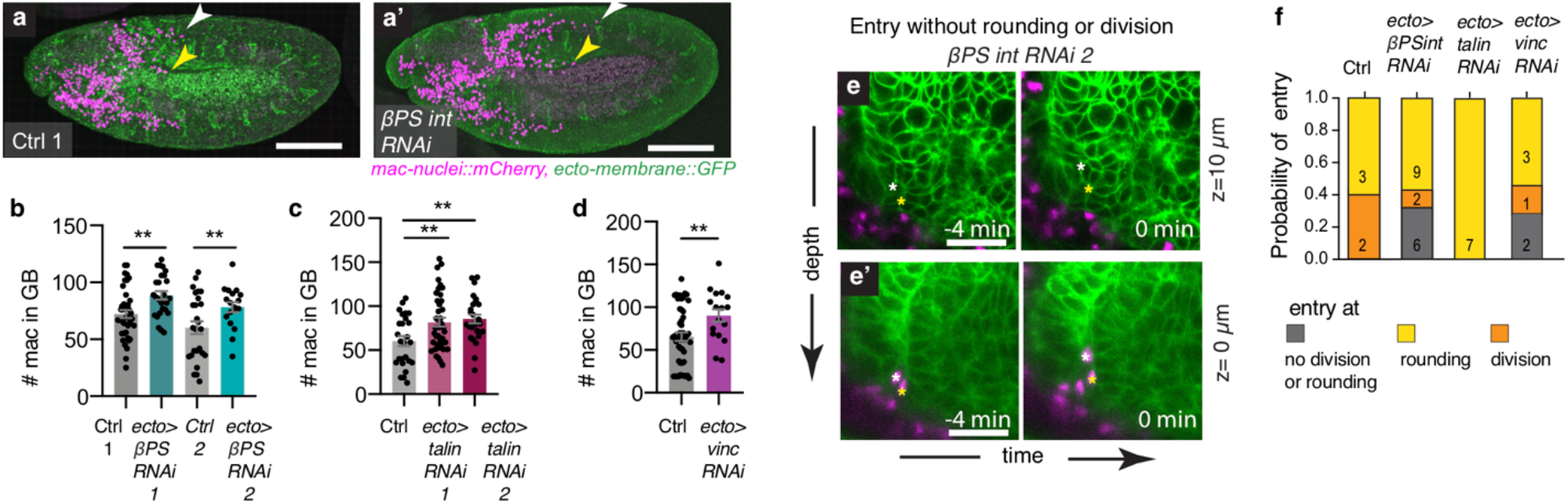
Reducing ectodermal FA components experimentally enables macrophage entry without mitotic rounding. **a,a’.** Maximum intensity projections of stage 12 embryos: control and ectodermal *βPS integrin RNAi*. **b-d,** Macrophage numbers inside the germband in embryos expressing (**b**) *βPS int RNAis*, (**c**) *talin RNAis* or (**d**) *vinculin RNAi* in the ectoderm compared to controls; means±s.e.m, (b)**p=0.0022, n=40,26; **p=0.0025, n=26,18 (c)**p=0.0020, **p=0.0039, n=26, 26, 26; (d) **p=0.0026, n=25, 20, unpaired t-tests. **e, e’,** Timelapse images of *ecto*>*βPS int RNAi* embryo in z-plane 10μm above macrophage (**e**) and in z-plane of the first macrophage (**e’**) entering without ectodermal cell rounding at the entry point. Nucleus labeled with a white star; the second macrophage labeled with a yellow star. **f.** Probability distribution of macrophage entry in embryos, sorted into different categories as indicated. Scale bars: 100μm (**a**); 20μm (**e**).

## Discussion

What controls cells’ ability to infiltrate into tissues is little understood, but has broad implications for development, cancer and immunology as all three processes require such movement. By using live imaging, focused genetic manipulations and quantitative analyses we demonstrate that macrophage invasion critically depends on tissue cell division. We identify the disassembly during mitosis of the focal adhesions that attach a cell to its surroundings as the key step by which division enables infiltration. Such disassembly has also been found for vertebrate cells^26,27^. Thus, our findings suggest that programmed or induced tissue division could permit the localization of vertebrate tissue resident immune cells, as well as those recruited during inflammation. Interestingly, surrounding tissue cell division can be triggered by macrophages themselves, through secretion of growth factors that also enable tissue repair^29^. Macrophages may thus effect their own tissue entry as well as that of other cells, particularly since macrophages have been identified as crucial partners for the tissue invasion of immune and cancer cells^30,31^. Moreover, tumor progression and metastases have been strongly linked to the level of infiltration by tumor associated macrophages^31,32^. At the same time, proliferative capacity remains a major indicator of tumor progression^33^. Our study suggests that these two factors can form a positive feedback loop: proliferation rates in solid tumors could control macrophage infiltration, which in turn can promote cancer cell dissemination. Indeed, a correlation has been observed, with higher proliferating tumors displaying higher macrophage infiltration^34–36^. Thus, our work should prompt an examination of how surrounding tissue cell division affects migratory cell behavior in a wide range of normal and disease contexts.

## Acknowledgements

We thank J. Friml, C. Guet, T. Hurd, M. Fendrych and members of the laboratory for comments on the manuscript; the Bioimaging Facility of IST Austria for excellent support and T. Lecuit, E. Hafen, R. Levayer and A. Martin for fly strains. This work was supported by a grant from the Austrian Science Fund FWF: Lise Meitner Fellowship M2379-B28 to M.A and D.S., and internal funding from IST Austria to D.S. and EMBL to S.D.R.

## Author contributions

M.A. and D.S. conceived the study and wrote the manuscript, with critical feedback from all authors. M.A. discovered the phenomena with valuable input from A.R. M.A. and D.S designed the experiments. M.A. performed and analyzed the experiments, with the help of A.G., M.V., F.V., A.A. and S.E. A.A. wrote the data analysis script. D.K and S.D.R hosted and guided M.A. during the optogenetic experiments.

---

**Fig. S1 related to Fig.1.**
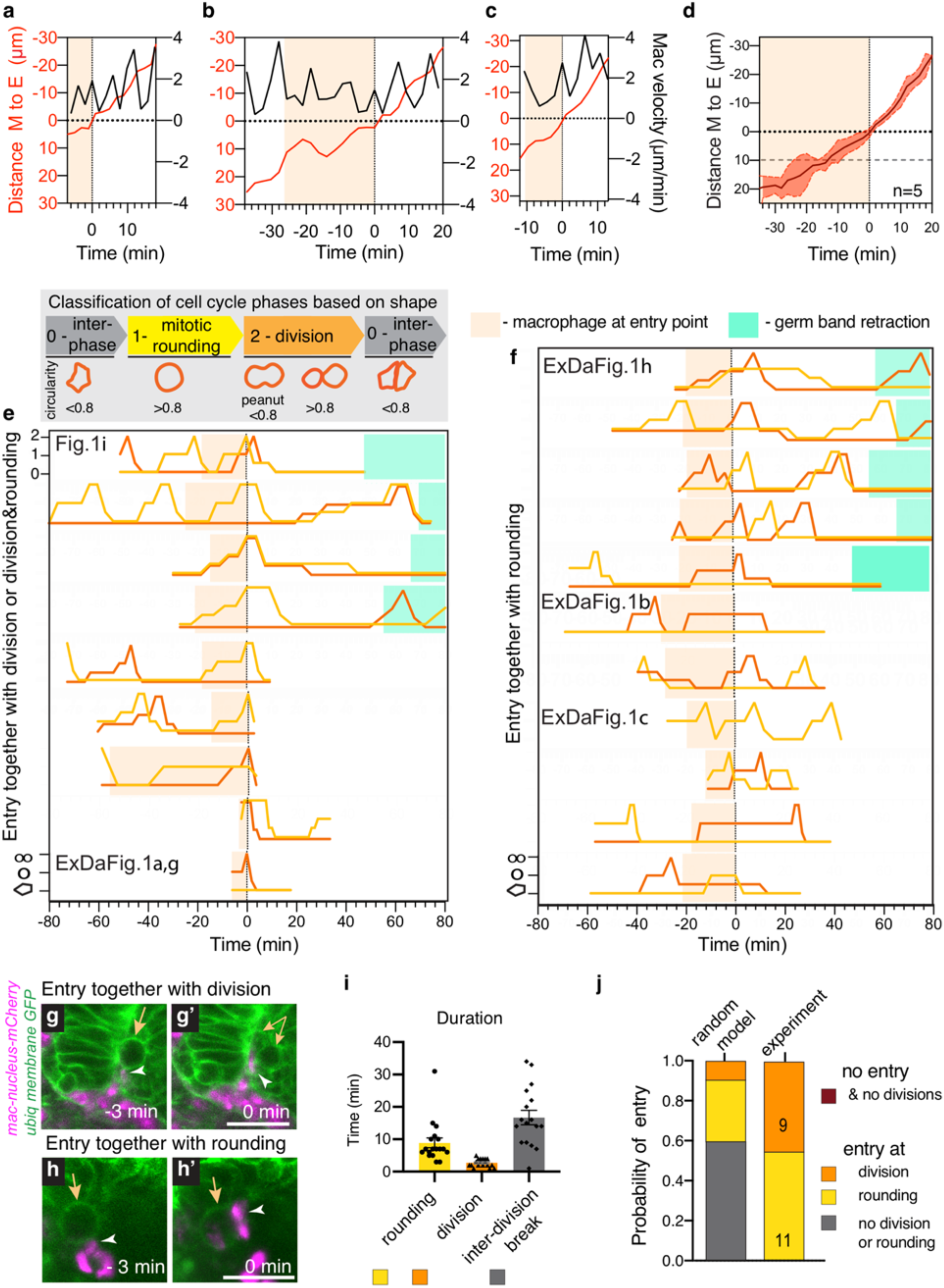
Correlation of macrophage entry and adjacent cellular mitosis is not due to frequent divisions at the entry point. **a-c,** Analysis of the first entering macrophage’s path and speed: the distance from the macrophage nucleus (M) to the entry point (E) is plotted over time (red line). Peach colored shading indicates that the macrophage nucleus is <10 μm from the entry point. **d,** Cumulative graph of ME distances; mean (solid line), ± s.d (shadowed colored area) (n=5 embryos). **e-f,** Division profiles, showing that each macrophage enters (**e**) alongside at least one dividing cell or (**f**) adjacent to at least one rounded cell. Bottom, middle and top positions on the *y*-axis indicate that a cell is in interphase, undergoing the mitotic rounding which starts at prophase, or undergoing division (in telophase & cytokinesis till daughter cells maintain rounded shape), as shown in the cell outlines to the lower left. Cell shape was assessed using the membrane markers. When the cell shape was not completely round (circularity >0.8, but has acute angles), the line is drawn between the lower and middle level. Peach colored shading represents the time period when the macrophage nucleus is <10 μm from the germband entry point. Green shading indicates when the germband is retracting. Time resolution: 1-4 min. The profiles whose speed/distance are shown in the stipulated Figures above are indicated. **g-h**, Time-lapse images of a macrophage entering as the adjacent cell (**g**) divides, or (**h**) rounds before division. Genotype: *Resille::GFP*, *DE-Cad::GFP*; *srpHemo-H2A::3xmCherry*. **i,** Quantification of the duration of the different cell cycle phases of the cells at the entry point calculated from the division profiles in panels **e-f**. In all embryos at most two ectodermal cells adjacent to the macrophage entry point undergo mitosis. Thus, the duration of the rounding or division phases is calculated as the uninterrupted time period when at least one of the two cells is undergoing the respective phase. The interdivision break duration is the time when neither of the two cells is rounded or dividing. Data are means± s.e.m. N=18, 18, 17 data points for rounding, division and interdivision break respectively from n=20 embryos. **j,** The probability of a macrophage entering together with ectodermal cell rounding, division or without either during the interdivision break. Left bar: the theoretical probability assuming that macrophages enter at random time points once they arrive at the entry point. The probability of entry during a particular phase is proportional to the average duration of each of three phases, presented in **i**. Right: the experimentally observed probability of entry for n=20 embryos (**e-f**). Note that in the experiment macrophages never enter during the interdivision break.

**Fig. S2 related to Fig.1.**
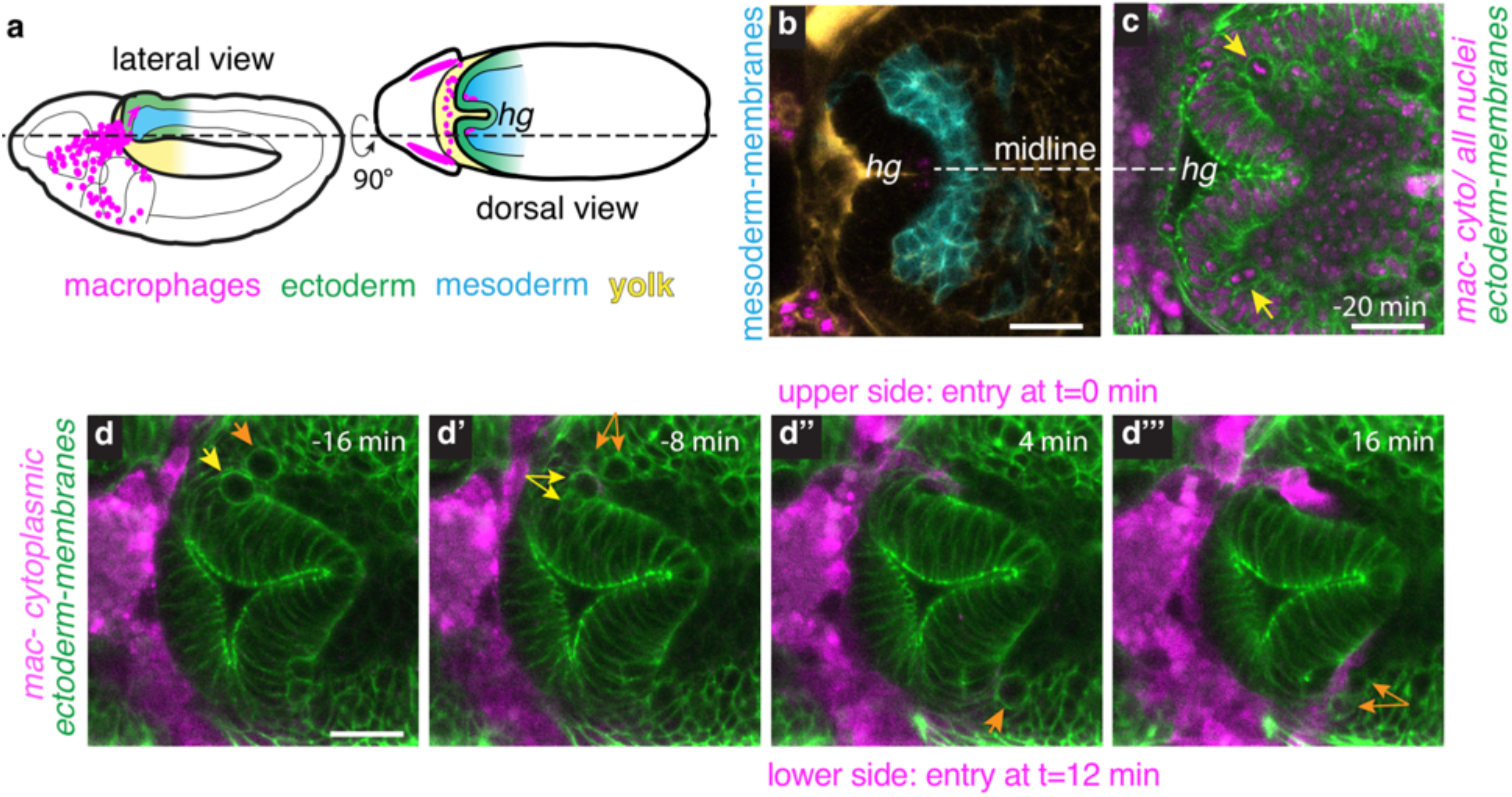
Macrophages enter symmetrically but not synchronously on both sides of the hindgut, due to temporally offset ectodermal cell divisions. **a,** Schematic of a lateral and dorsal view of the embryo showing that macrophages enter into the germband on both sides of the hindgut (hg). **b,** Confocal image of the dorsal view of the ecto-meso interface close to the entry point with mesodermal membranes labeled in blue as in (**a**)**. c,** Confocal image of the dorsal view of the ecto-meso interface close to the entry point with nuclei of all cells labeled (magenta) along with ectodermal membranes (green) (*ubi-H2A::RFP*; *knock-in DE-Cad::GFP*). Mitotic cells at both entry points (yellow arrows) are located symmetrically with respect to the midline, but divisions are not synchronized. **d,** Dorsal view time-lapse images of macrophage entry. (**d-d’’**) On the upper side macrophages enter just after the division of two ectodermal cells. (**d’’’**) 12 minutes later on the lower side macrophages enter after the division of one ectodermal cell. Macrophages labeled by the cytoplasmic marker *srpHemo-3xmCherry*, ectodermal membranes by *knock-in DE-Cad::GFP*. Scale bars in all panels: 20μm.

**Fig. S3.**
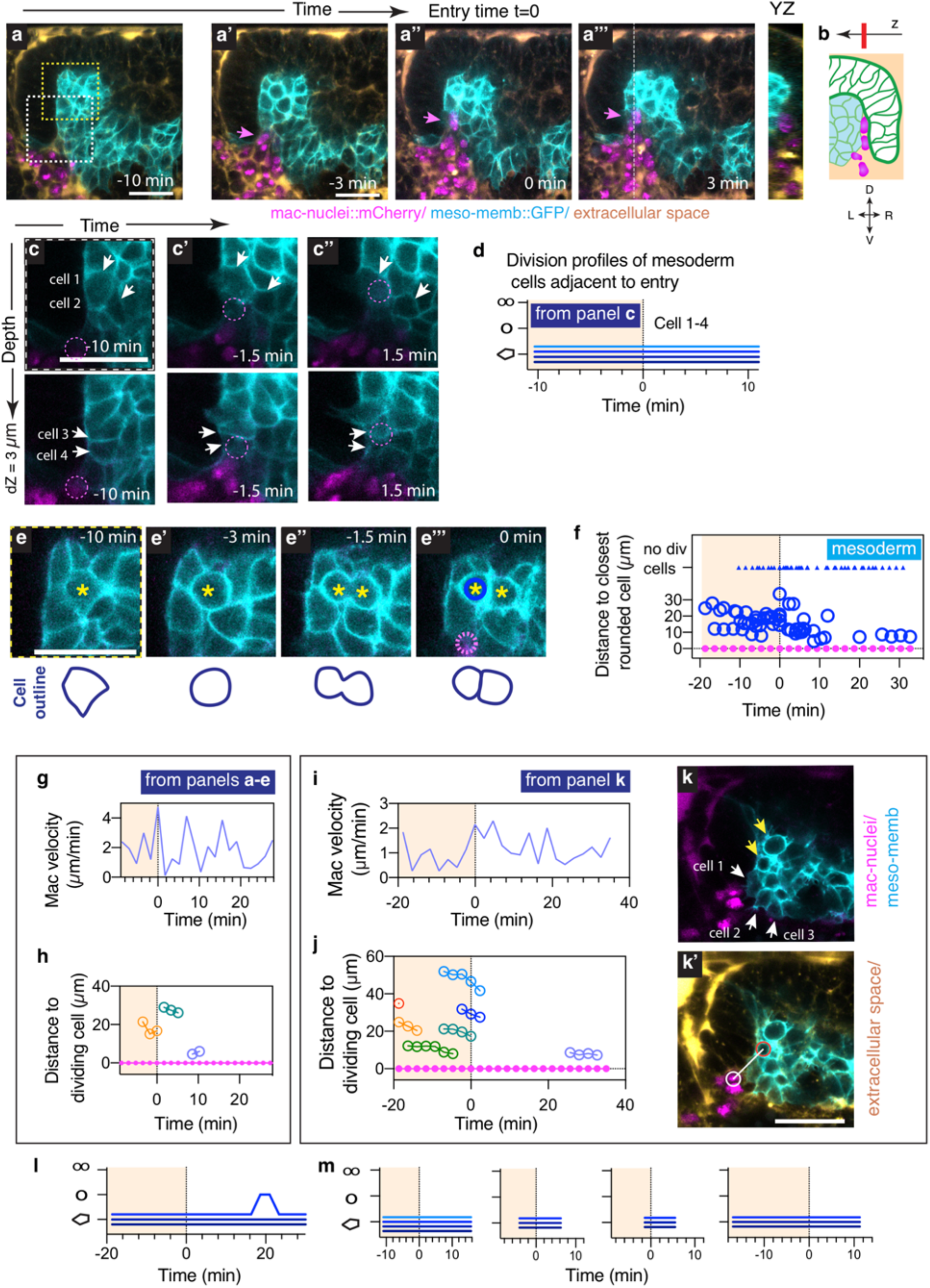
Macrophage entry is not correlated with mitosis of mesodermal cells. **a-a’’’,** Confocal timelapse imaging of WT embryo. Mesodermal cell membranes visualized by *VT45198-GAL4*>*10xUAS-IVS-myr::GFP*; macrophage nuclei labeled with *srpHemo-H2A::3xmCherry*, and extracellular space visualized by injection of dextran-Alexa-fluor647, 10,000MW. Macrophage entry time denoted by t=0 min. Max intensity projection of 3 slices with dZ=1μm. Dashed white line in (**a’’’,** left panel) shows the position of the plane of the orthogonal view (right). **b,** Scheme of the orthogonal view in **a’’’** with a red line depicting the position of the cross section shown in **a-a’’’**. **c,** Zoom of dashed white box in **a**. Upper row of images: mesodermal cells (cells 1, 2) immediately adjacent to the entering macrophage, whose nucleus in located 3 μm above these images at a position depicted by the dashed magenta circle. Lower row of images: 3 μm deeper mesodermal cells (cells 3, 4) which eventually touch the macrophage during entry. **d,** Division profiles of cells 1-4 from panel **c.** None of these cells is rounding or dividing during macrophage entry. **e,** Zoom of dashed yellow box in **a**: a mesodermal cell undergoing mitotic rounding and division before macrophage entry and at a distance from the entry point. **f**, Distance from the first macrophage to the center of the closest round mesodermal cell over time. Data points are shown as circles with the average radius of these round cells demonstrating that the macrophage (magenta) is not close to any round mesodermal cell at the time of entry. Image **e’’’**: example of the macrophage nucleus (magenta circle) and the closest round cell (blue circle) at entry. Triangles show cases in which there is no rounded mesodermal cell closer than 40μm from the macrophage nucleus. Cumulative graph from n=10 embryos. **g,** Nuclear velocity of the first entering macrophage from the embryo presented in panels **a-e.** **h,** Corresponding graph of the distance from the first macrophage to the center of the round mesodermal cells within a 40μm radius around the macrophage. Linked one-color circles represent the same cell in consecutive frames. Data points are shown as circles with the average radius of a round cell to demonstrate that the macrophage is not near any round cell at the time of entry. Magenta dots indicate the location of the macrophage nucleus. **i-j,** Graphs as in **g-h** of data from embryo presented in panel **k**. **k,** Confocal image of the entry region, showing the first macrophage 20 minutes before entry and all mesodermal cells dividing at that time point (two yellow arrows) within a 50μm radius around the macrophage. Membranes of mesodermal cells visualized by *VT45198-GAL4*>*10xUAS-IVS-myr::GFP*; macrophage nuclei labeled with *srpHemo-H2A::3xmCherry*. **k’**, The same image with extracellular space visualized by injection of dextran-Alexa-fluor 647 (yellow), and a schematic of the distance between the macrophage and the closest dividing mesodermal cell. **l-m,** Division profiles of mesodermal cells located at the ecto-meso interface at the entry site shows that none of these cells is rounding or dividing during macrophage entry. **l** follows cells 1-3 marked on panel **k**. **m** shows similarly located cells from 4 other embryos.

**Fig. S4 related to Fig.2.**
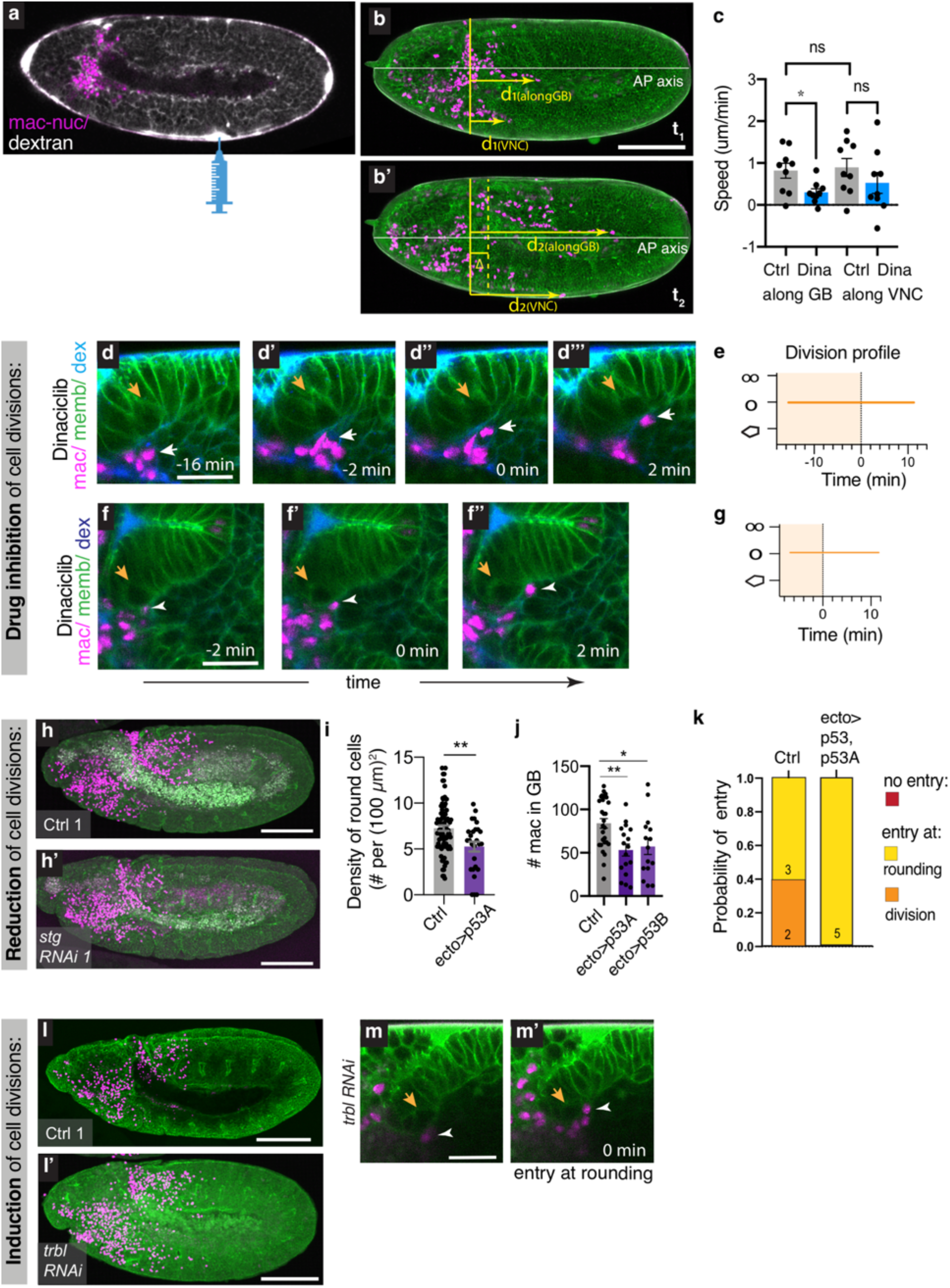
Inhibition of ectodermal cell divisions impairs macrophage entry, and induction enhances it. **a,** The location of the injections is shown by the syringe. To show the diffusion speed from this site throughout the extracellular space of the embryo, we imaged 10 min after injecting dextran-Alexa-fluor 647, 10,000MW (white). **b-b’,** Scheme of the distances *d_1_* and *d_2_* measured at times *t_1_* and *t_2_* to quantify the average directional speed of macrophages. Δ is the shift of the germband (GB) edge due to retraction. Horizontal white line shows anterior to posterior (AP) axis. **c,** Quantification of macrophage speed after dinaciclib injection indicates that macrophages are motile and move in an AP direction both along the edge of the germband and ventrally along the ventral nerve cord (vnc). Speed is relative to surrounding tissues and calculated from time-lapse images starting before GBR and ending at ~40% germband retraction (GBR) using the following equations: V_alongGB_=(d_2(alongGB)_-d_1(alongGB)_-Δ)/(t_2_-t_1_); V_along vnc_=(d_2(vnc)_-d_1(vnc)_+Δ)/(t_2_-t_1_), where Δ is the shift of the GB edge due to retraction. A Δ μm posterior retraction by the germband results in an anterior movement by the vnc of the same distance, hence the opposite sign for Δ in the two equations. *p=0.0465, one-way ANOVA with Tukey. **d-d”‘,** Lateral confocal time-lapse zoomed images of macrophage entry in an embryo injected with dinaciclib and dextran, showing the juxtaposition of the entering macrophage nucleus (white arrow) and a rounded ectodermal cell (orange arrow). **e,** Division profile of the rounded cell in (**d**) over time showing that it did not divide in a 30 min time period, but maintained its round shape. **f-f”.** Dorsal confocal time-lapse zoomed images of another dinaciclib and dextran-injected embryo showing the first macrophage (white arrowhead) entering the germband next to a rounded ectodermal cell at the entry site (orange arrow) that (**g**) shows no division for 20 min. **h-h’** Representative confocal images of Stage 12 fixed embryos showing that fewer macrophages infiltrated into the germband in embryos that express *stg RNAi* in the ectoderm (*ecto*>*stg RNAi*) than in the control embryos. **i,** Density of the rounded cells in the germband tissue just before macrophage entry in the control (n=10 embryos, z=4-7 per embryo, sample size=83) and *ecto*>*UASp53,p53A* (n=6 embryos, z=4-7 per embryo, sample size=27) showing reduced division density upon overexpression of p53 (**p<0.01), unpaired t-test. **j,** Quantification of macrophage numbers inside the germband reveal a tissue invasion defect upon overexpressing p53 in the ectoderm: *ecto*>*UASp53,p53A* (n=17, **p<0.01) and *ecto*>*UASp53,p53B* (n=15, *p<0.05) compared to control (n=39); one-way ANOVA test with Tukey. **k,** Probability distribution of macrophage entry: macrophages enter into the germband together with ectodermal cell rounding (light orange) in *ecto*>*UASp53,p53A* (n=4; control n=5). **l-l’,** Representative confocal images of Stage 12 fixed embryos showing that more macrophages infiltrated the germband in embryos that express *tbl RNAi* in the ectoderm (*ecto*>*tbl RNAi*) than in control embryos. **m-m’.** Time lapse images of an *ecto*>*trbl RNAi* embryo showing macrophage entry neighboring a mitotically rounded ectodermal cell. Scale bars: 100μm (**h-h’**, **l-l’**), 10μm (**m-m’**) Genotype: *e22cGAL4*, *Resille::GFP*, *DE-Cad::GFP/RNAi or UASp53,p53B*; *srpHemo-H2A::3xmCherry*/+.

**Fig. S5.**
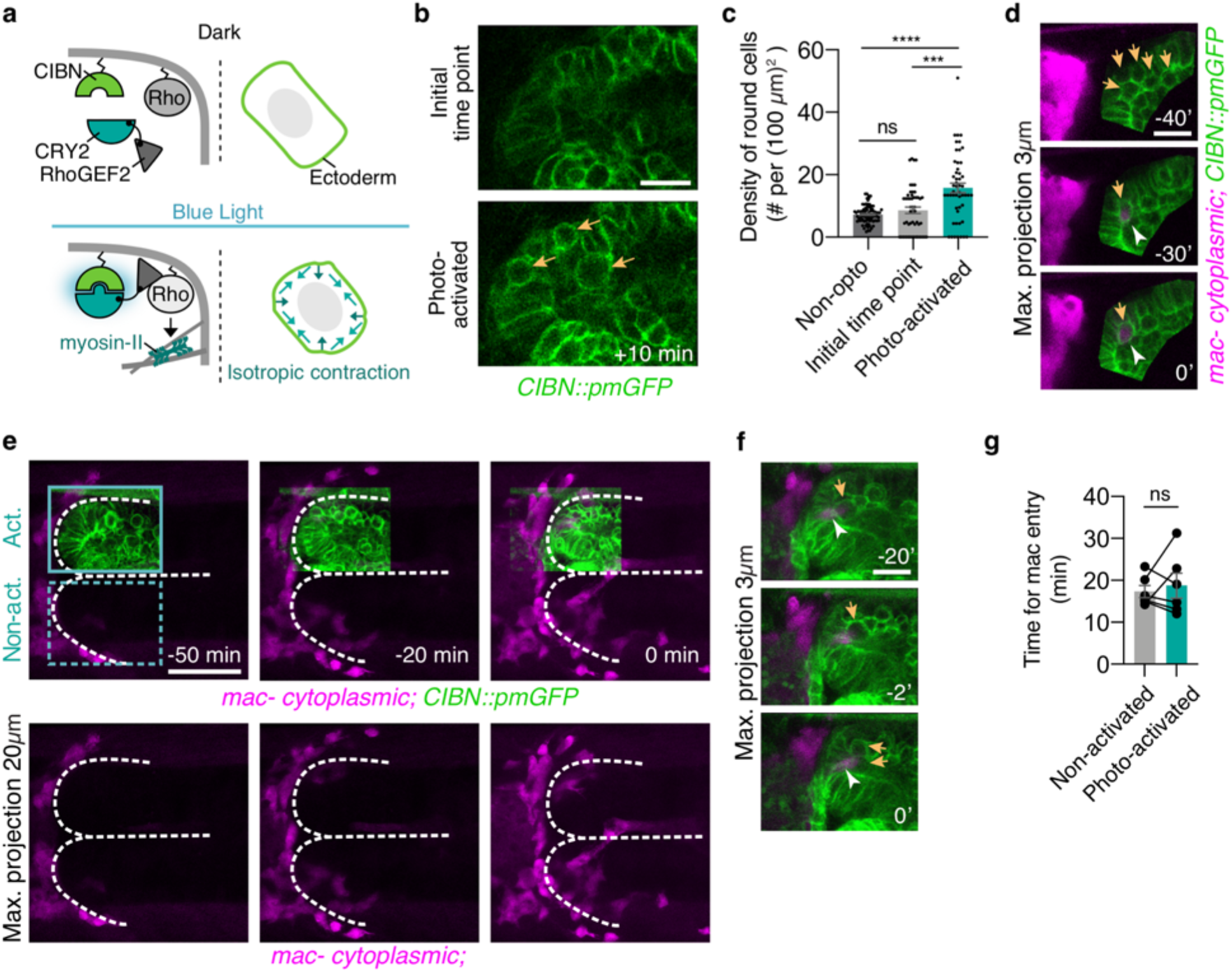
Optogenetic activation of cortical contractility in ectodermal cells and subsequent rounding of cells at the entry site doesn’t enhance macrophage invasion. **a,** Schematic of the optogenetic system expressed in ectodermal cells for the induction of cortical actomyosin contractility by the activation of RhoGEF2 with the blue light. It is composed of the photosensitive cryptochrome 2 (CRY2) fused to the catalytic domain of RhoGEF2, and its GFP-tagged binding partner CIBN anchored at the plasma membrane. In the dark, RhoGEF2-CRY2 is present in the cytoplasm and inactive (top). Blue light illumination triggers a conformational change in CRY2 enforcing its interaction with CIBN, which thus causes the translocation of RhoGEF2-CRY2 to the plasma membrane, where it activates endogenous Rho1 signaling (bottom) and induces myosin-mediated contractility. **b,** Representative two-photon images of the activated region of the ectoderm at the initial time point revealing a round cell density similar to the non-activated ectoderm (upper panel); and after 5 cycles of photo-activation (lower panel) showing induced rounding of ectodermal cells at the germband edge beside the entry point (orange arrows). The density of round cells increases ~2 times after 5 cycles (10 min) of activation. **c**, Rounded cell density in the germband tissue just before macrophage entry in the control (n=83, N=10 embryos, z=5-6 z-planes per embryo); at the initial time point and in the illuminated region after 10 cycles of photo-activation (n=51 each data set, N=8 embryos, z=5-6 z-planes per embryo), ***P=0.0002, Wilcoxon paired test; ns P=0.93, ****P<0.0001 one-way ANOVA with Kruskal–Wallis test followed by post hoc Dunn’s test. **d**, Representative maximum intensity projections of an activated region of the ectoderm and macrophages, showing that cell rounding at the entry point induced by activation precedes macrophage appearance at the entry point (upper image) and that a rounded cell is present continuously at the entry point from 40 min before the entry time onwards. **e**, Maximum intensity projections of dorsal view embryos, visualizing the 50×40μm region activated by blue light (green) at −050 min and macrophages (magenta) over time; macrophage entry in the activated half occurs at time=0 min. White dashed line outlines the edge of the germband and its midline. **f,** Images of the activated region in the embryo presented in (**e**), showing that a rounded cell is present at the entry point at the time when the macrophage first reaches it. The macrophage enters together with division of this cell. **g,** Quantification of the time macrophages reside at the entry point before they enter into the non-activated region or activated region, showing no effect of the induced ectodermal cortex contraction (n=6 embryos); ns: p=0.616, paired t-test. Scale bars: 20μm (**b,d,f**), 50μm (**e**).

**Fig. S6 related to Fig.3.**
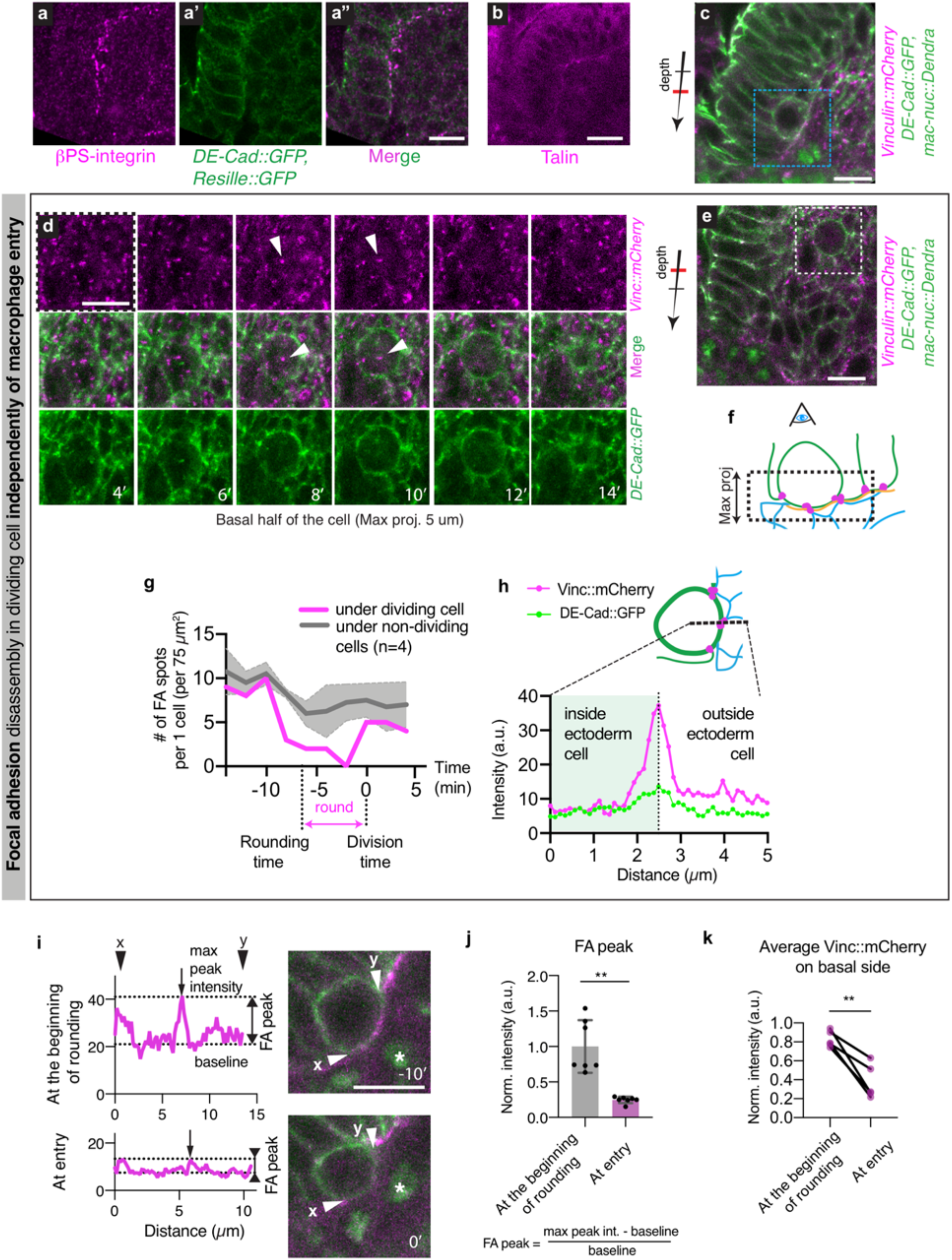
The ectoderm-mesoderm interface is sealed by Integrin-containing focal adhesions (FA) which gradually disappear during ectodermal mitotic rounding, independently of macrophage presence. **a-a”,** Confocal images of the ectoderm-mesoderm interface in an embryo, (**a**) immunostained for βPS-integrin to show FAs, (**a’**) with cell membranes labeled by DE-Cad::GFP, Resille::GFP, and (**a”**) merged, showing that the ecto-meso interface contains Integrin puncta. Slice position corresponds to the scheme in Fig.1**b**, left panel. **b,** Live confocal images of Talin::mCherry at the ectoderm-mesoderm interface in the edge of the germband. **c**, Confocal image of an overview of the ecto-meso interface, blue square region zoomed in Fig. 3d. **d**, Maximum intensity projection of the dotted white square region in (**e**) showing an example of FA disassembly in the dividing cell seen dorsally. Upper row: Vinculin::mCherry maximum intensity projection showing all FAs. Middle row: merge. Lower row: Membranes visualized by DE-Cad::GFP. Notice the single FA spot in the middle of the basal side of the round cell at time 8’, and disassembled by time 12’. Scheme of the maximum projection orientation is shown in (**f**). **e,** Overview image showing the position of a rounding cell far from the entry point and thus from signals potentially emitted by macrophages. Dotted white square region is zoomed in panel (**d**). **f,** Scheme of the orientation of the maximum intensity projection shown in panel (**d**). Mesoderm shown in blue, the rounding ectodermal cell and its neighbors in green, Vinculin::mCherry in magenta, and the ECM in orange. **g.** Quantification of the number of FAs spots per individual ectodermal cell from images in panel (**e**). Gray curve shows the average number of FAs under a non-dividing ectodermal cell (per the area corresponding to one cell). Pink curve shows the average number of FAs under a dividing cell. Solid gray line represents mean and shadowed area ± s.d. (n=1 dividing cell and n=5 non-dividing cells). On x-axis 0=time of division. Time when mitotic rounding begins is indicated. **h,** Intensity profiles orthogonal to the cell membrane of Vinculin::mCherry (pink) and DE-Cad::GFP (green) in the rounded cell show that the peaks of both channels coincide. Schematic above shows the position of the line ROI used for the profiles. **i,** Examples of Vinculin::mCherry profiles along the basal side of the rounding cell shown on the right, defining peak and baseline intensities. **j,** Vinculin::mCherry peak intensity on the basal side of mitotically rounding cells at the entry point at the beginning of rounding and at the time of macrophage entry. The intensity amplitude of the FA peak was quantified as the difference between the maximum intensity of the peak and the baseline intensity, normalized by the baseline intensity: (peak-base)/base; illustrated in (**i**). Values are further normalized to the average of the intensity amplitude at the beginning of rounding (n=7 embryos, **P=0.002, paired t-test). **k,** Average intensity of Vinculin::mCherry on the basal side of mitotically rounding cells at the entry point at the beginning of rounding and at the time of macrophage entry (n=5), normalized by basal intensity before rounding. **p=0.0053, paired t-test. Scale bars: 10μm in (**a**-**e, i**).

**Fig. S7 related to Fig.4.**
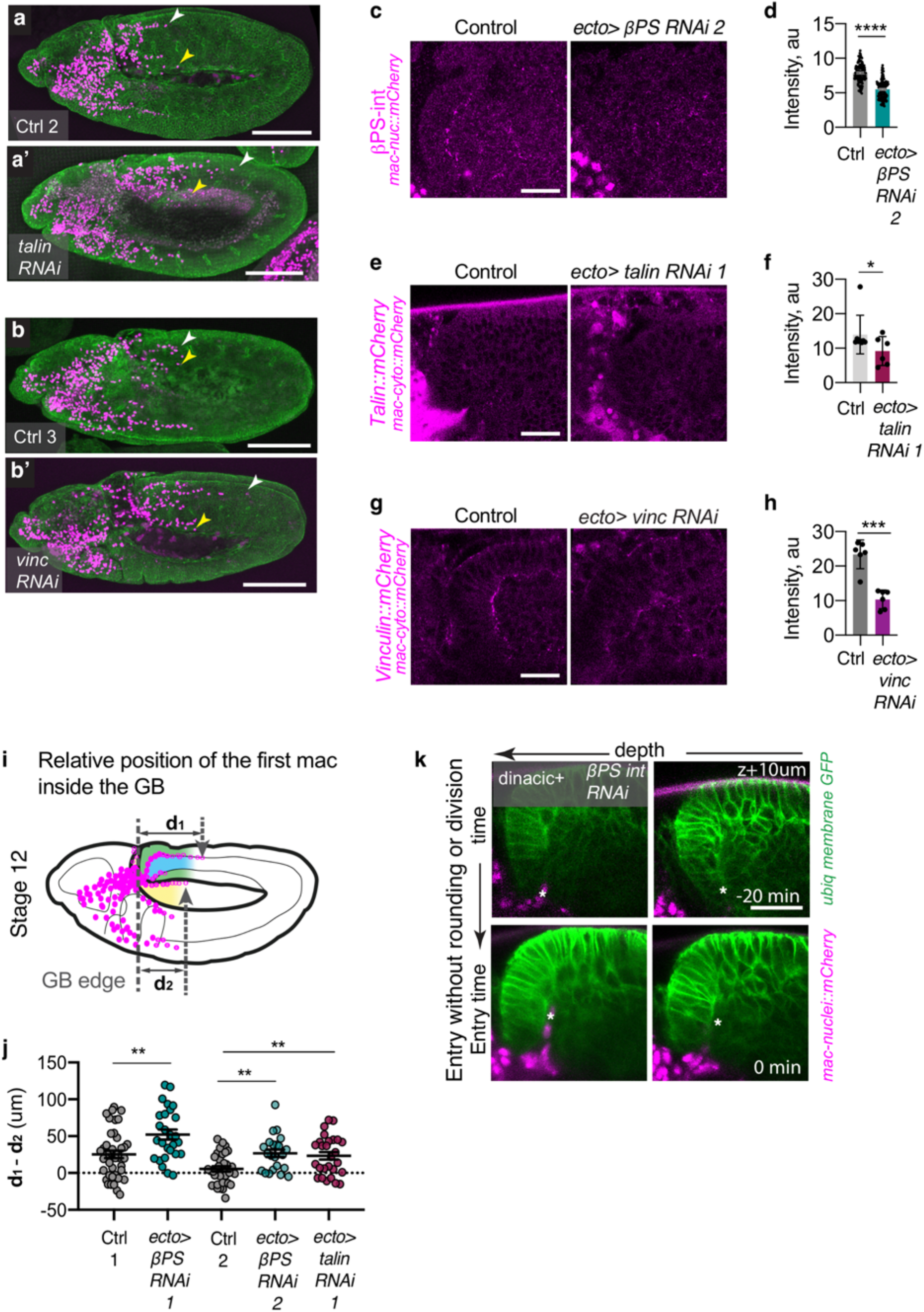
Reducing FA components in the ectoderm accelerates macrophage entry and enables macrophage entry without ectodermal mitotic rounding. **a-b’,** Representative confocal images of Stage 12 fixed embryos demonstrated that compared to the control (**a,b**) more macrophages infiltrated into the germband when adhesion components were reduced by (**a’**) ectodermally expressed *talin RNAi* or (**b’**) *vinculin RNAi*. **c,** Representative confocal images and (**d**) quantification of the fluorescence intensity of fixed control and *ecto*>*βPS int RNAi* embryos treated with fluorescently labeled anti-*β*PS integrin antibody. **e,** Representative confocal images and (**f**) quantification of the fluorescence intensity of live embryos expressing Talin::mCherry alone (control) or along with *ecto*>*talin RNAi*. **g,** Representative confocal images and (**h**) quantification of the fluorescence intensity of live embryos expressing Vinculin::mCherry alone (control) or along with *ecto*>*vinc RNAi*. **i,** Scheme of the distances *d_1_* and *d_2_* measured for the examination of macrophage invasion efficiency in (**j**) quantifying macrophage migration inside the dense germband tissue compared to the non-invasive route along the germband edge. **j,** Plotted is the difference between the distance migrated from the GB end by the macrophage farthest inside the germband (*d_1_*) and the distance migrated by the farthest macrophage outside the germband (*d_2_*). Upon the reduction of adhesion components in the ectoderm, macrophages migrate farther inside the germband than in the controls. Control 1 (n=40), *βPS int RNAi* 1=VDRC103704 (n=26), **p<0.01; control 2 (n=26), *βPSint RNAi* 2=BL33642 (n=18) **p<0.01; *talin RNAi 2*=BL33913 (n=26), p<0.01; *vinc RNAi*=BL25965 (n=17), **p<0.01 by one-way ANOVA with Tukey’s test. **k,** Timelapse images of an *ecto*>*βPS int RNAi* embryo, injected with dinaciclib, in which macrophages enter without ectodermal cell rounding at the entry point. White star indicates the first entering macrophage nucleus. Scale bars: 100μm in (**a-b’**); 20μm in (**k**).

**Fig. S8.**
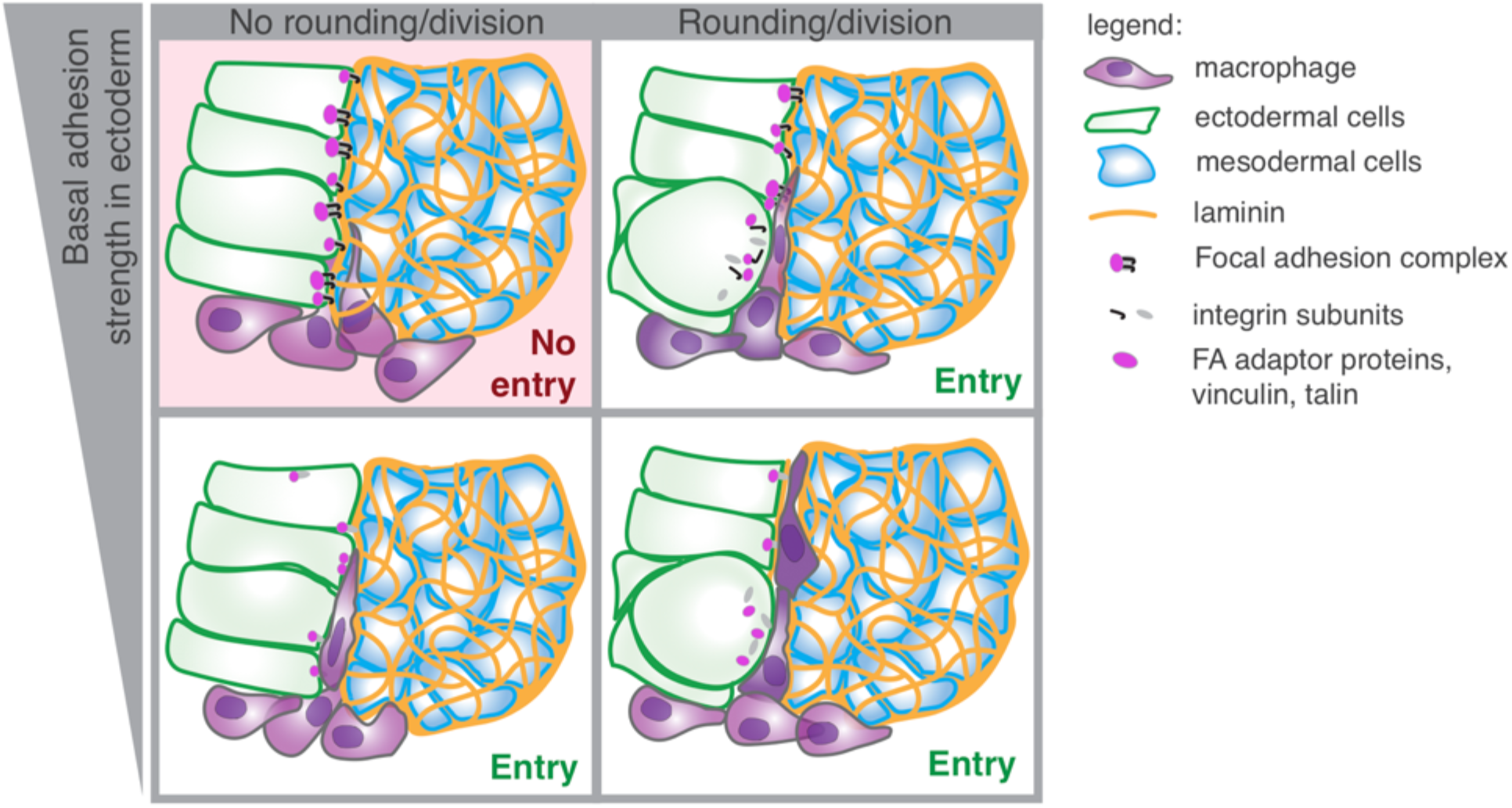
Graphical abstract. 1) In wild type embryos macrophages cannot invade into the confined space between the ectoderm and mesoderm if no ectodermal cells are rounding up for mitosis at the entry site. This is due to the firm attachment between the ectoderm and mesoderm on the edge of the germband tissue which does not allow the macrophage nucleus to penetrate between the adhesion foci. 2) Ectodermal cell division is thus required for the first macrophage to invade due to the focal adhesion disassembly that occurs during mitotic rounding. 3) A knockdown of adhesion components is sufficient to enable macrophages to enter independently of divisions: macrophage entry can occur with or without a mitotic ectodermal cell adjacent to the entering macrophage.

## Methods

### Fly strains and genetics

Flies were raised at room temperature on food bought from IMBA (Vienna, Austria) containing agar, cornmeal, and molasses with the addition of 1.5% Nipagin. For embryo collections, adults were placed in cages in a Sanyo 555 MIR-153 incubator at 29°C and 65% humidity; embryos were collected on apple juice plates prepared in house and containing sugar, agar and Nipagin, supplemented with dry yeast from Lesaffre (Marcq, France) on the plate surface. Embryo collections for RNA interference experiments were done at 29°C to optimize expression under GAL4 driver control. Embryos were collected after 7.5 h at 29°C for fixation and after 4-4.5 h for live imaging.

The Berkeley Drosophila Genome Project’s *in situ* database^37,38,39^ and the FlyBase^40^ database of *Drosophila* genes were consulted to examine the embryonic expression of genes and to find stocks.

Fly lines used in the study are listed below, complete genotypes of which are provided in Supplementary Methods. *srpHemo-H2A::3xmCherry* and *srpHemo::3xmCherry* lines have been previously described^41^. The knock-in *DECad::GFP* line in which GFP is fused to the C terminus of Cadherin and knocked into the endogenous locus was provided by Y. Hong^42^; the ubiquitous membrane marker *Resille::GFP* is a gift from A. Martin^43^; *Ubi-H2A::mRFP* from L.Ringrose^44^; *UAS-myc::p53A*, *UAS-myc::p53B* from E. Hafen; *UAS-cycD*, *UAS-cdk4* from R. Levayer and *Vinculin*:*mCherry*, *DECad::GFP* from T. Lecuit^45^. The following lines were obtained from the Bloomington Stock Centre: *e22c-GAL4*^46^; *10XUAS-IVS-myr::GFP*^47^; *UAS-p53*; *Talin::mCherry MiMIC line*^48^; TRiP lines^49^ *UAS-RNAi* for *stg(string)*, *mys*(*βPS-integrin*), *rhea*(*talin*), *vinculin*. The following lines were obtained from the Vienna Drosophila Resource Center (VDRC), Vienna, Austria: *VT45198-GAL4*^50^, *UAS-RNAi of mys* (*βPS-integrin*), *trbl*(*tribbles*) lines^51^. Optogenetic line *w[*]*; *P{w*+, *UASp-CIBN::pmGFP}*; *P{w*+, *UASp-RhoGEF2-CRY2}* is described in ^25^.

The *P{w*+ *srpHemo-mCerulean::H2B::Dendra}* line was constructed as follows. The srpHemo promoter fragment was inserted at the Stu1 restriction site of the PCasper4 plasmid^52^ using the infusion kit from Clontech and the following infusion primer pair: AGGTCGACCTCGAGGCCTAAATTTTGATGTTTTTAAATAGTCTTATCAGCAATGGCAA AACGTTAACTCGAGGCCTTATGGGATCCGTGCTGGGGTAGTGC

The resulting pC4-srpHemo plasmid was cut with Xba1 and a fragment from the mCerulean-H2B::Dendra plasmid^53^ (a gift from the C.P. Heisenberg lab) was inserted downstream of the srpHemo promoter fragment using the infusion kit from Clontech with the following primer pair: CGAGGTCGACTCTAGATCCCATCGATATGGGCTG ATCTGGATCCTCTAGACATGCGTTTAAACCCGGG

Injection into flies at random locations was conducted as described by Gyoergy et al^41^.

### Embryo fixation and immunohistochemistry

Embryos collected on apple juice plates for 7.5 h at 29°C and afterwards kept at 4°C for up to 48 h were incubated in 50% Chlorox (DanClorix) for 5 min and washed. Embryos were fixed with 17% formaldehyde/heptane (ThermoFisher Scientific, Waltham, MA, USA) for 20 min followed by methanol devitellinization. For *β*PS-integrin antibody staining, embryos were fixed with 4.0% paraformaldehyde from a freshly prepared, frozen and then thawed stock along with heptane for 20 min followed by methanol devitellinization as described previously^54^. Antibodies used: mouse anti-Integrin *β*PS ^55^ (DSHB CF.6G11, 1:25), incubated overnight at 4°C. Afterwards, embryos were washed in BBT for 2 h, incubated with secondary Alexa fluor 633 labelled antibodies at a dilution of 1:500 (Thermo Fisher Scientific, Waltham, Massachusetts, USA) at RT for 2 h, and washed again for 2 h. After immunolabeling embryos were incubated in Vectashield (Vector Laboratories, Burlingame, USA) o/n at 4°C, mounted on a slide and imaged.

### Imaging of fixed embryos

Embryos containing an ectodermal GFP membrane marker and a nuclear mCherry macrophage marker and found in a lateral orientation were imaged in two channels through the whole embryo with a Zeiss Inverted LSM800 or upright LSM900 Confocal Microscope using a Plain-Apochromat 20X/0.8 Air Objective with XY-resolution 0.62 μm and Z-resolution of 3 μm (~70 μm total stack). Antibody-stained embryos were imaged with an upright LSM900 Confocal Microscope using a Plain-Apochromat 40X/1.3 Oil Objective with a XY-resolution of 0.2 μm and a Z-resolution of 0.5 μm with constant excitation intensities.

### Live imaging

Embryos collected on apple juice plates for 4 h at 29°C were incubated in 50% Chlorox (DanClorix) for 5 min and washed. Stage 10-11 embryos were selected based on autofluorescence in blue light using a fluorescent stereomicroscope and glued to an18×18mm high precision coverslip (Marienfeld Laboratory 699 Glassware, No. 1.5H) in the lateral or dorsal orientation, placed on silica beads for 5 min for dehydration and then covered with a droplet of halocarbon oil 200 (Sigma) and an oxygen permeable membrane (YSI). A 100×100 μm size or smaller anterior dorsolateral region of the embryo was imaged immediately with a Zeiss Inverted LSM800 or upright LSM900 Confocal Microscope using a Plain-Apochromat 40X/1.3 Oil Objective and a temperature control unit set to 29°C, at an XY-resolution of 0.15 μm, Z-resolution of 1 μm and resulting time resolution of 90 to 360 s depending on the size of the area imaged. Channels were imaged sequentially and excitation intensities were adjusted to the tissue penetration depth.

For injections, embryos were dechorionated in 50% Chlorox for 2 min, washed, and mounted in a lateral orientation, then dehydrated by placing on silica beads for 15-20 mins. Embryos were covered with a droplet of halocarbon oil 200 (Sigma) and injected with either dinaciclib (Selleckchem, Cat# S2768) resuspended to 500 μM in DMSO or with the addition of 0.1 mg/ml 10 KDa Dextran Alexa Fluor 647 (10,000, Invitrogen) or undiluted DMSO. Injections were performed using a Femto Jet Injectman (Eppendorf) with Femto tips II (Eppendorf) into the perivitelline space on the ventral side of Stage 10 embryos, in which macrophages have just approached the germband edge. Imaging was performed 15 min after injection. Five to six live embryos mounted on one coverslip were imaged sequentially with a Zeiss Inverted LSM800 using a Plain-Apochromat 20X/0.8 Air Objective with XY-resolution of 0.62 μm and Z-resolution of 3 μm (~40 μm stack comprising half of the embryo), time resolution 10-12 min.

### Live imaging in optogenetic experiments

Flies expressing the optogenetic module were kept in the dark. Stage 11 embryos were selected in halocarbon oil using a standard stereomicroscope illuminated with a red-light emitting LED lamp. Embryos were dechorionated with 100% sodium hypochlorite for 1 min, washed with distilled water and mounted onto a 35-mm glass-bottom dish (MatTek corporation) in PBS with their dorsal side facing the coverslip.

Optogenetic experiments were performed on a commercial Zeiss LSM 780 NLO confocal microscope (Carl Zeiss) equipped with a tunable (690–1,040 nm) femtosecond (140 fs) pulsed laser (Chameleon; Coherent, Inc.), which operates at a repetition rate of 80 MHz, and using a C-Apochromat 40x/1.20 W Corr FCS M27 water immersion objective (Carl Zeiss) at 20°C. A Deep Amber lighting filter (Cabledelight, Ltd) was used to filter bright field illumination while locating the samples under the microscope. Zen Black software (Carl Zeiss) in combination with the Pipeline Constructor macro (Politi et al, 2018) was used to operate the microscope. The mCherry reporter signal expressed in macrophages was recorded and simultaneously transmission light was collected using the HeNe-laser at 561-nm excitation in an image stack spanning ~200 μm in the x-y dimension and a total z-stack size of ~35 μm at 1 μm intervals. Selective optogenetic activation of RhoGEF2-CRY2 in one half of the ectodermal tissue without photoactivating the other half on the opposite side of the midline was achieved using two-photon excitation at λ=950 nm. By reference to images acquired with transmitted light collected using the 561 nm laser, a ~50×40×20 μm spanning volume in which cells were photo-activated was defined and illuminated with a z-interval of 0.5 μm. Photo-activation of the sample volume was done with a laser power of 18 mW, a pixel size of 0.42 μm, a pixel dwell time of 2.5 μs and comprising 3 consecutive iterations. The time to complete an entire photo-activation cycle was ~70 s. An initial (pre-activation) mCherry z-stack was acquired, followed by alternating cycles of photo-activation and mCherry reporter acquisition.

### Image data analysis

Images were processed and analyzed using Fiji^56^ (https://fiji.sc/), Bitplane Imaris 9.3 and homemade scripts (Python).

#### Tracking and velocity calculation

Macrophage nuclei were tracked in laterally oriented embryos using TrackMate^57^ plugin in Fiji (https://imagej.net/TrackMate) with manual correction. A circle was drawn around the first entering macrophage and tracked forward and backward in time. The intersection point of the ecto/meso interface, basal ectodermal membrane and the edge of the mesoderm was defined as an entry point. Its position was verified in the entry frame, where it is situated immediately above the entering macrophage nucleus. The entry point was marked manually in TrackMate and tracked. The distance between the first macrophage nucleus and the entry point, d(t), was calculated from their XYZ positions saved in the tracking files. d(t) values are declared positive before entry, and negative after entry for the convenience of velocity calculations. Macrophage velocity relative to the entry point was calculated as follows:

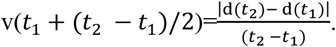

Distances between the macrophage nucleus and the closest rounded mesoderm cell in Fig. S3 were measured similarly using XYZ positions saved in tracking files. Visualization of 3D rendered videos was performed in Bitplane Imaris 9.3, with manual segmentation of round cells and automated segmentation of macrophage nuclei using Surface function.

#### Division profiles

Cell shapes of ectodermal or mesodermal cells adjacent to the entry point were analyzed manually by drawing cell outlines on top of the membrane channel in Fiji. Values 0, 1, 2 were assigned to image frames based on cell circularity (=4*pi*area/perimeter^2) in their central cross section: 0 for *circ* <0.8; 1 for *circ* >0.8 (round cell without acute angles); 2 for peanut-shape dividing cell and two smaller round daughter cells with *circ* >0.8, which replace a bigger round cell. A value of 0.5 was assigned to the cell with *circ* >0.8 having acute angles or straight edges, which mostly occur in the daughter cells after division. Quantification of the difference in time between the closest division or mitotic rounding event and entry (Fig. 1j) uses the closest timepoint to entry time having values 2 or 1 respectively, in either of the two cells’ division profiles.

#### Density of rounded cells

Live images of the germband’s anterior end at 20 to 10 min before macrophage entry were used for quantification. In each z-stack freehand regions of interest (ROIs) were drawn in Fiji, comprising the anterior part of the ectoderm up to 100 μm from the germband edge, in 5-7 z-slices comprising 20 μm in total. The number of round cells with *circularity*>0.8 was counted manually inside the ROIs and divided by ROI area to calculate round cell density.

#### Image and data analysis of optogenetic experiments

Image stacks of the macrophage-mCherry channel were compiled with the GFP channel of the activated region (first z-stack of every activation cycle) using Fiji. The density of round cells was calculated similarly as described above by defining an ROI inside the activated region, which contained the outermost layer of ectodermal cells. The area of the ROI was calculated in Fiji and the number of round cells inside the ROI was determined manually based on the shape of the membrane labeling CIBN:pmGFP signal (*circularity*>0.8) in 5-6 z-slices with a 2 μm interval. The presence of mitotically round cells at the entry point in the activated region was assessed manually by analyzing cell shapes adjacent to the entering macrophage as described above.

The time for macrophage entry was calculated as follows. A maximum intensity projection of the mCherry channel in a dorsal view was performed in Fiji. Time 1, when the first macrophage marked with the cytoplasmic *srpHemo-3xmCherry* was first seen to move under the germband to a position ~15 μm from the GB edge in the posterior direction, determined the time when the macrophage first reaches the entry point. From this position macrophages, which are entering into the germband tissue, move in a dorsal direction. Thus, Time 2, corresponding to macrophage entry, was determined by an abrupt two-fold increase in the mCherry intensity, corresponding to the movement of the first macrophage towards the surface of the embryo during entry. The time for macrophage entry was calculated as the difference between Time 2 and Time 1.

#### Macrophage counting in the germband

The membrane marker channel was used to measure the position of the germband front end relative to the embryo length and to identify the edge of the germband tissue. Fixed embryos of Stages 11-12 were identified by having a stomodeal invagination and germband retraction away from the anterior between 29-40% of the embryo length. The number of macrophage nuclei within the germband was calculated manually in Fiji. In injected live embryos the number of macrophage nuclei within the germband was calculated at 30-31% of germband retraction.

*The number of adhesion spots per cell* was calculated by outlining cell membranes on the basal surface of the ectodermal cells in a maximum projection of 5 μm (4 z-slices), thresholding the FA signal and manually calculating the number of FA spots inside outlined areas, then dividing by the area.

#### Analysis of fluorescence intensities

Quantification of peak fluorescence intensities of basal focal adhesions was performed on raw images by manually outlining the basal side of the ectodermal cell (line of 5 pixels width) and applying the Plot Profile function in Fiji. The maximum intensity value in the middle region of the profile was taken as a peak intensity and the average of the profile outside the peak as a baseline. The amplitude of the peak intensity was calculated as: (maximum peak intensity–baseline)/baseline. To calculate average Vinculin::mCherry intensities on the basal side fluorescence intensity values were averaged along the line profiles (line of 5 pixels width) and between profiles of the same cell in all z-slices and normalized to the cytosolic signal. Mean intensity within the ectoderm for *β*PS-integrin staining, Talin::mCherry and Vinculin::mCherry to assess RNAi effect was assessed within ROIs inside the ectoderm tissue, normalized to the background signal outside embryo. 10-15 ROIs from 8-10 embryos were used for the analysis.

### Statistical analysis and repeatability

All statistical analyses were performed with GraphPad Prism v9.0.1. The normality test was performed by D’Agostino and Pearson or Shapiro-Wilk methods, and the appropriate test to evaluate statistical differences between the means was chosen based on the distribution as indicated in Figure legends: two-tailed unpaired t-test, paired t-test or Mann-Whitney test to compare two groups and one-way ANOVA with Tukey’s test for multiple comparisons, indicating p-values *P < 0.05, **P < 0.01, ***P < 0.001 and ****P < 0.0001. In Fig. 1j the box plot extends from the 25th to the 75th percentiles, with the middle line showing the mean, and the whiskers indicating the minimum to maximum values.

All measurements and analysis were performed in 4-50 embryos. Images presented in figures are representative of separate biological replicates repeated at least 3 and up to 20 times. Stills from confocal movies shown in figures are representative of independent experiments repeated at least 4 and up to 20 times as indicated in Figure legends.

### Exact genotype of *Drosophila* lines used in Figures

**Fig.1 c-j, Fig. S1:** *Resille::line was constructed as followsGFP*, *DE-Cad::GFP*; *P{w*+ *srpHemo-H2A::3xmCherry}*.

**Fig. S2b:** *P{10XUAS-IVS-myr::GFP}attp40*; *VT45198-GAL4*, *P{w*+ *srpHemo-H2A::3xmCherry};* **c:** *P{Ubi-H2A::mRFP,w*^+^*}; DE-Cad::GFP, P{w*+ *srpHemo-3xmCherry};* **d:** *DE-Cad::GFP, P{w*+ *srpHemo-3xmCherry}*. **Fig. S3:** *P{10XUAS-IVS-myr::GFP}attp40*; *P{VT045198-GAL4}, P{w*+ *srpHemo-H2A::3xmCherry}*.

**Fig. 2b-g & Fig. S4a-g:** *Resille::GFP, DE-Cad::GFP; P{w*+ *srpHemo-H2A::3xmCherry}*.

**Fig. 2 h-I & Fig. S4h: Control 1:** *yw-; P{en2.4-GAL4}e22c, Resille::GFP, DE-Cad::GFP*/+; *P{w*+ *srpHemo-H2A::3xmCherry}/ P{CaryP}attP2* (made from BDSC8622). **ecto**>**stg RNAi 1:** *yw-; P{en2.4-GAL4}e22c, Resille::GFP, DE-Cad::GFP/*+; *srpHemo-H2A::3xmCherry/P{TRiP.JF03235}attP2*. **Control 2:** *yw-; P{en2.4-GAL4}e22c, Resille::GFP, DE-Cad::GFP /P{CaryP}attP40; srpHemo-H2A::3xmCherry*/+ *(made from BDSC36304)*. **ecto**>**stg RNAi 2:** *yw-; P{en2.4-GAL4}e22c, Resille::GFP, DE-Cad::GFP/ P{TRiP.GL00513}attP40; P{w*+ *srpHemo-H2A::3xmCherry}*/+.

**Fig. S4i-k: Control:** *w-; P{en2.4-GAL4}e22c, Resille::GFP, DE-Cad::GFP*/+; *P{w*+ *srpHemo-H2A::3xmCherry}*/+. **ecto**>**p53**: *P{en2.4-GAL4}e22c, Resille::GFP, DE-Cad::GFP/P{w[*+*mC]=GUS-p53}2.1; P{w*+ *srpHemo-H2A::3xmCherry}/ UAS-myc-p53A or UAS-myc-p53B* (made from BDSC6584).

**Fig. 2m-o & Fig. S4l,m: Control 1:***yw-; P{en2.4-GAL4}e22c, Resille::GFP, DE-Cad::GFP/P{attP,y[*+*],w[3]*+ *(made from VDRC 60100GD); P{w*+ *srpHemo-H2A::3xmCherry}*/+], **ecto**>**trbl RNAi 1:** *yw-; P{en2.4-GAL4}e22c, Resille::GFP, DE-Cad::GFP/UAS-trbl RNAi VDRC106774KK; P{w*+ *srpHemo-H2A::3xmCherry*/+*}*. **Control 2:** *w-; P{en2.4-GAL4}e22c, Resille::GFP, DE-Cad::GFP*/+; *P{w*+ *srpHemo-H2A::3xmCherry}*/+.

**ecto**>**trbl RNAi 2:** *yw-; P{en2.4-GAL4}e22c, Resille::GFP, DE-Cad::GFP*/+; *P{w*+ *srpHemo-H2A::3xmCherry}/ UAS-trbl RNAi VDRC22114GD*. **ecto**>**cycD, cdk4**: *w-; P{en2.4-GAL4}e22c, Resille::GFP, DE-Cad::GFP/UAS-cycD, UAS-cdk4; P{w*+ *srpHemo-H2A::3xmCherry}*/+.

**Fig. S5:** *w[*]; P{w*+, *UASp-CIBN::pmGFP}/P{en2.4-GAL4}e22c, P{w*+ *srpHemo-3xmCherry}; P{w*+, *UASp-RhoGEF2-CRY2}*/+.

**Fig. 3 & Fig. S6:** *w; pFlyFos Vinc::mCherry, DE-Cad::GFP; P{w*+ *srpHemo-mCerulean::H2B::Dendra}*.

**Fig. S6b:** Talin::mCherry*: y[1] w[*]; Mi{PT-mCh.0}rhea[MI00296-mCh.0]/TM6B, Tb[1]* (BDSC39648).

**Fig. 4a,b,e,f & Fig. S7c,d,i-k: Control 1:** *y w-; P{en2.4-GAL4}e22c, Resille::GFP, DE-Cad::GFP /P{attP,y[*+*],w[3’](VDRC 60100GD); P{w*+ *srpHemo-H2A::3xmCherry}*/+.

**ecto**>β**PS-integrin RNAi 1**: *yw-/w-; P{en2.4-GAL4}e22c, Resille::GFP, DE-Cad::GFP/UAS-mys RNAi VDRC*103704/KK*; P{w*+ *srpHemo-H2A::3xmCherry}*/+.

**Control 2:** *yw-/w-; P{en2.4-GAL4}e22c, Resille::GFP, DE-Cad::GFP*/+; *P{w*+ *srpHemo-H2A::3xmCherry}/P{CaryP}attP2* (made from BDSC8622).

**ecto**>β**PS-integrin RNAi** 2: *yw-/yv; P{en2.4-GAL4}e22c, Resille::GFP, DE-Cad::GFP*/+; *P{w*+ *srpHemo-H2A::3xmCherry}/ P{y[*+*t7.7] v[*+*t1.8]= TRiP.HMS00043}attP2* (made from BDSC33642).

**Fig. 4c,f & Fig. S7I,j: Control:** *yw-/w-; P{en2.4-GAL4}e22c, Resille::GFP, DE-Cad::GFP*/+; *P{w*+ *srpHemo-H2A::3xmCherry}/ P{CaryP}attP2* (made from BDSC8622).

**ecto**>**talin RNAi 1:** *y w-/y sc v; P{en2.4-GAL4}e22c, Resille::GFP, DE-Cad::GFP*/+; *P{w*+ *srpHemo-H2A::3xmCherry}/P{y[*+*t7.7] v[*+*t1.8]=.HMS00856}attP2* (made from BDSC33913).

**ecto**>**talin RNAi 2:** *y w-/y v; P{en2.4-GAL4}e22c, Resille::GFP, DE-Cad::GFP*/+; *P{w*+ *srpHemo-H2A::3xmCherry}/ P{y[*+*t7.7] v[*+*t1.8]=TRiP.HM05161}attP2* (made from BDSC28950).

**Fig. 4d,f: Control:** *y w-/w-; P{en2.4-GAL4}e22c, Resille::GFP, DE-Cad::GFP*/+; *P{w*+ *srpHemo-H2A::3xmCherry}/ P{CaryP}attP2* (made from BDSC8622).

**ecto**>**vinc RNAi:** *y w-/y v; P{en2.4-GAL4}e22c, Resille::GFP, DE-Cad::GFP*/+; *P{w*+ *srpHemo-H2A::3xmCherry}/ P{y[*+*t7.7] v[*+*t1.8]=TRiP.JF01985}attP2* (made from BDSC25965).

**Fig. S7e,f: Control:** female *P{en2.4-GAL4}e22c, Resille::GFP, DE-Cad::GFP, P{w*+ *srpHemo::3xmCherry}; Talin::mCherry/TM3, P{GAL4-twi.G}2.3, P{UAS-2xEGFP} AH2.3, Sb[1] Ser[1] (made with BL6663*) x male *yw*-;+; *P{CaryP}attP2*.

**ecto**>**talin RNAi 1:** female *P{en2.4-GAL4}e22c, Resille::GFP, DE-Cad::GFP, P{w*+ *srpHemo::3xmCherry; Talin::mCherry/TM3, P{GAL4-twi.G}2.3, P{UAS-2xEGFP} AH2.3, Sb[1] Ser[1] (made with BDSC6663*) x male *yw*-;+; *P{y[*+*t7.7] v[*+*t1.8]=TRiP.HMS00856}attP2*.

**Fig. S7g,h: Control:** *P{en2.4-GAL4}e22c, P{10XUAS-IVS::myr::GFP}attp1, P{w*+ *srpHemo::3xmCherry}, vinculin::mCherry/vinculin::mCherry*;+/*TM3, P{GAL4-twi.G}2.3, P{UAS-2xEGFP} AH2.3, Sb[1] Ser[1] (made with BL6663*)

**ecto**>**vinc RNAi:** *P{en2.4-GAL4}e22c, P{10XUAS-IVS::myr::GFP}attp1, P{w*+ *srpHemo-3xmCherry}, vinculin::mCherry/vinculin::mCherry*;+/*P{y[*+*t7.7] v[*+*t1.8]=TRiP.JF01985}attP2*.

## References

1. Friedl, P. & Weigelin, B. Interstitial leukocyte migration and immune function. Nature Immunology 9, 960–969 (2008).

2. Friedl, P. & Alexander, S. Cancer invasion and the microenvironment: Plasticity and reciprocity. Cell 147, 992–1009 (2011).

3. Kulesa, P. M. & Gammill, L. S. Neural crest migration: Patterns, phases and signals. Developmental Biology 344, 566–568 (2010).

4. Kameritsch, P. & Renkawitz, J. Principles of Leukocyte Migration Strategies. Trends in Cell Biology 30, 818–832 (2020).

5. Charras, G. & Sahai, E. Physical influences of the extracellular environment on cell migration. Nat. Rev. Mol. Cell Biol. 15, 813–824 (2014).

6. Nia, H. T., Munn, L. L. & Jain, R. K. Physical traits of cancer. Science (New York, N.Y.) 370, (2020).

7. Yamada, K. M. & Sixt, M. Mechanisms of 3D cell migration. Nature Reviews Molecular Cell Biology 20, 738–752 (2019).

8. Siekhaus, D., Haesemeyer, M., Moffitt, O. & Lehmann, R. RhoL controls invasion and Rap1 localization during immune cell transmigration in Drosophila. Nat. Cell Biol. 12, 605–610 (2010).

9. Ratheesh, A. et al. Drosophila TNF Modulates Tissue Tension in the Embryo to Facilitate Macrophage Invasive Migration. Dev. Cell 45, 331–346.e7 (2018).

10. Achilleos, A. & Trainor, P. A. Neural crest stem cells: Discovery, properties and potential for therapy. Cell Research 22, 288–304 (2012).

11. Kierdorf, K., Prinz, M., Geissmann, F. & Gomez Perdiguero, E. Development and function of tissue resident macrophages in mice. Seminars in Immunology 27, 369–378 (2015).

12. Park, C. O. & Kupper, T. S. The emerging role of resident memory T cells in protective immunity and inflammatory disease. Nature Medicine 21, 688–697 (2015).

13. Casano, A. M., Albert, M. & Peri, F. Developmental Apoptosis Mediates Entry and Positioning of Microglia in the Zebrafish Brain. Cell Rep. 16, 897–906 (2016).

14. Luster, A. D., Alon, R. & von Andrian, U. H. Immune cell migration in inflammation: Present and future therapeutic targets. Nature Immunology 6, 1182–1190 (2005).

15. Meirson, T., Gil-Henn, H. & Samson, A. O. Invasion and metastasis: the elusive hallmark of cancer. Oncogene 39, 2024–2026 (2020).

16. Benias, P. C. et al. Structure and distribution of an unrecognized interstitium in human tissues. Sci. Rep. 8, (2018).

17. Stewart, T. A., Hughes, K., Hume, D. A. & Davis, F. M. Developmental Stage-Specific Distribution of Macrophages in Mouse Mammary Gland. Front. Cell Dev. Biol. 7, 250 (2019).

18. Munro, D. A. D. & Hughes, J. The origins and functions of tissue-resident macrophages in kidney development. Frontiers in Physiology 8, 837 (2017).

19. Epelman, S., Lavine, K. J. & Randolph, G. J. Origin and Functions of Tissue Macrophages. Immunity 41, 21–35 (2014).

20. Ratheesh, A., Belyaeva, V. & Siekhaus, D. Drosophila immune cell migration and adhesion during embryonic development and larval immune responses. Curr. Opin. Cell Biol. 36, 71–79 (2015).

21. Valoskova, K. et al. A conserved major facilitator superfamily member orchestrates a subset of O-glycosylation to aid macrophage tissue invasion. Elife 8, (2019).

22. Belyaeva, V. et al. Cortical actin properties controlled by Drosophila Fos aid macrophage infiltration against surrounding tissue resistance. bioRxiv 2020.09.18.301481 (2020). doi:10.1101/2020.09.18.301481

23. Emtenani, S. et al. A genetic program boosts mitochondrial function to power macrophage tissue invasion. bioRxiv 2021.02.18.431643 (2021). doi:10.1101/2021.02.18.431643

24. Taubenberger, A. V, Baum, B. & Matthews, H. K. The Mechanics of Mitotic Cell Rounding. Mech. Mitotic Cell Rounding. Front. Cell Dev. Biol 8, 687 (2020).

25. Izquierdo, E., Quinkler, T. & De Renzis, S. Guided morphogenesis through optogenetic activation of Rho signalling during early Drosophila embryogenesis. Nat. Commun. 9, 1–13 (2018).

26. Dix, C. L. et al. The Role of Mitotic Cell-Substrate Adhesion Re-modeling in Animal Cell Division. Dev. Cell 45, 132–145.e3 (2018).

27. Lock, J. G. et al. Reticular adhesions are a distinct class of cell-matrix adhesions that mediate attachment during mitosis. Nat. Cell Biol. 20, 1290–1302 (2018).

28. Klapholz, B. & Brown, N. H. Talin - The master of integrin adhesions. Journal of Cell Science 130, 2435–2446 (2017).

29. Wynn, T. A. & Vannella, K. M. Macrophages in Tissue Repair, Regeneration, and Fibrosis. Immunity 44, 450–462 (2016).

30. Stolp, B. et al. Salivary gland macrophages and tissue-resident CD8+ T cells cooperate for homeostatic organ surveillance. Sci. Immunol. 5, (2020).

31. Cassetta, L. & Pollard, J. W. Tumor-associated macrophages. Curr. Biol. 30, R246–R248 (2020).

32. Aras, S. & Raza Zaidi, M. TAMeless traitors: Macrophages in cancer progression and metastasis. British Journal of Cancer 117, 1583–1591 (2017).

33. Van Diest, P. J., Brugal, G. & Baak, J. P. A. Proliferation markers in tumours: Interpretation and clinical value. Journal of Clinical Pathology 51, 716–724 (1998).

34. Garvin, S. et al. Differences in intra-tumoral macrophage infiltration and radiotherapy response among intrinsic subtypes in pT1-T2 breast cancers treated with breastconserving surgery. Virchows Arch. 475, 151–162 (2019).

35. Hao, N. B. et al. Macrophages in tumor microenvironments and the progression of tumors. Clinical and Developmental Immunology 2012, (2012).

36. Qiu, S. Q. et al. Tumor-associated macrophages in breast cancer: Innocent bystander or important player? Cancer Treatment Reviews 70, 178–189 (2018).

## References

37. Hammonds, A. S. et al. Spatial expression of transcription factors in Drosophila embryonic organ development. Genome Biol. 14, (2013).

38. Tomancak, P. et al. Systematic determination of patterns of gene expression during Drosophila embryogenesis. Genome Biol. 3, (2002).

39. Tomancak, P. et al. Global analysis of patterns of gene expression during Drosophila embryogenesis. Genome Biol. 8, (2007).

40. Larkin, A. et al. FlyBase: Updates to the Drosophila melanogaster knowledge base. Nucleic Acids Res. 49, D899–D907 (2021).

41. Gyoergy, A. et al. Tools Allowing Independent Visualization and Genetic Manipulation of Drosophilamelanogaster Macrophages and Surrounding Tissues. G3 (Bethesda). g3.300452.2017 (2018). doi:10.1534/g3.117.300452

42. Huang, J., Zhou, W., Dong, W., Watson, A. M. & Hong, Y. Directed, efficient, and versatile modifications of the Drosophila genome by genomic engineering. Proc. Natl. Acad. Sci. U. S. A. 106, 8284–8289 (2009).

43. Morin, X., Daneman, R., Zavortink, M. & Chia, W. A protein trap strategy to detect GFP-tagged proteins expressed from their endogenous loci in Drosophila. Proc. Natl. Acad. Sci. U. S. A. 98, 15050–15055 (2001).

44. Fonseca, J. P. et al. In vivo Polycomb kinetics and mitotic chromatin binding distinguish stem cells from differentiated cells. Genes Dev. 26, 857–871 (2012).

45. Kale, G. R. et al. Distinct contributions of tensile and shear stress on E-cadherin levels during morphogenesis. Nat. Commun. 9, 1–16 (2018).

46. Lawrence, P. A., Bodmer, R. & Vincent, J. P. Segmental patterning of heart precursors in Drosophila. Development 121, 4303–4308 (1995).

47. Pfeiffer, B. D., Truman, J. W. & Rubin, G. M. Using translational enhancers to increase transgene expression in Drosophila. Proc. Natl. Acad. Sci. U. S. A. 109, 6626–6631 (2012).

48. Venken, K. J. T. et al. MiMIC: A highly versatile transposon insertion resource for engineering Drosophila melanogaster genes. Nat. Methods 8, 737–747 (2011).

49. Perkins, L. A. et al. The Transgenic RNAi Project at Harvard Medical School: Resources and Validation. Genetics 201, 843–852 (2015).

50. Kvon, E. Z. et al. Genome-scale functional characterization of Drosophila developmental enhancers in vivo. Nature 512, 91–95 (2014).

51. Dietzl, G. et al. A genome-wide transgenic RNAi library for conditional gene inactivation in Drosophila. Nature 448, 151–156 (2007).

52. Thummel, C. S. & Pirrotta, V. Technical notes: new pCasper P-element vectors. D. I. S. 71, 150 (1992).

53. Dempsey, W. P., Fraser, S. E. & Pantazis, P. PhOTO Zebrafish: A Transgenic Resource for In Vivo Lineage Tracing during Development and Regeneration. PLoS One 7, e32888 (2012).

54. Zhang, L. & Ward IV, R. E. Distinct tissue distributions and subcellular localizations of differently phosphorylated forms of the myosin regulatory light chain in Drosophila. Gene Expr. Patterns 11, 93–104 (2011).

55. Brower, D. L., Wilcox, M., Piovant, M., Smith, R. J. & Reger, L. A. Related cell-surface antigens expressed with positional specificity in Drosophila imaginal discs. Proc. Natl. Acad. Sci. U. S. A. 81, 7485–7489 (1984).

56. Schindelin, J. et al. Fiji: an open-source platform for biological-image analysis. Nat. Methods 9, 676–682 (2012).

57. Tinevez, J.-Y. et al. TrackMate: An open and extensible platform for single-particle tracking. Methods 115, 80–90 (2017).

