## Supplementary Video Legends for "Cell division in tissues enables macrophage infiltration"

**Video 1.** The first macrophage enters the germband tissue adjacent to two dividing ectodermal cells. Left: macrophage nuclei labeled with *srpHemo-H2A::3xmCherry* (magenta). Middle: z-plane of the entering macrophage, corresponds to stills in **Fig. 1e.** Right: z=+6µm plane, corresponds to stills in **Fig. 1f.** Membranes labeled by Resille::GFP and DE-Cad::GFP. Scale bar: 20µm.

**Video 2.** Inhibition of cell division blocks entry. In embryos injected with dinaciclib to inhibit cell division macrophages enter into the germband tissue at the normal entry location and only when a round ectodermal cell is present at the entry site. Macrophage nuclei *srpHemo-H2A::3xmCherry* (magenta), membranes Resille::GFP and DE-Cad::GFP (green), dextran as a marker of dinaciclib injection (blue). Corresponds to **Fig. 2f.** Scale bar: 20µm.

**Video 3.** Correlation between entry and ectodermal division holds when the frequency of division is reduced in the ectoderm. Embryos shown express *stg* RNAi in the ectoderm. Left: Macrophages do not enter if no ectodermal cells are dividing at the germband edge. Corresponds to **Fig. 2j.** Right: Macrophages enter only if at least one ectodermal cell is dividing at the germband edge. Corresponds to **Fig. 2k.** Both lateral view. Macrophage nuclei *srpHemo-H2A::3xmCherry* (magenta), membranes Resille::GFP and DE-Cad::GFP (green). Scale bars: 20µm.

**Video 4.** Correlation between entry and ectodermal division holds when the frequency of division is increased in the ectoderm. Embryos shown express *trbl* RNAi in the ectoderm. Macrophages enter only if at least one ectodermal cell is dividing at the germband edge. Lateral view. Macrophage nuclei *srpHemo-H2A::3xmCherry* (magenta), membranes Resille::GFP and DE-Cad::GFP (green). Corresponds to Extended Data **Fig. 4m.** Scale bar: 20µm.

**Video 5.** Ectodermal focal adhesions disassemble during mitosis, leaving one central attachment which then also vanishes. This ectodermal cell is far from the macrophage entry point. Top view. Labeling: Vinculin::mCherry (magenta), DE-Cad::GFP (green). Corresponds to Extended Data **Fig. 6d.** Scale bar: 5µm.

**Video 6.** Macrophages enter when the last focal adhesion of the ectodermal cell disassembles during mitosis. At the entry point ectodermal focal adhesions disassemble after the cell rounds up during mitosis, leaving one central attachment, which diminishes (orange arrow). Lateral view. Entering macrophage nuclei labeled with Dendra (green traced circles, white arrow). Ectodermal labeling: Vinculin::mCherry (magenta), DE-Cad::GFP (green). Corresponds to **Fig. 3d.** Scale bar: 10µm.

**Video 7.** Reduction of the focal adhesion component, ß-integrin, in the ectoderm enables macrophage entry without mitotic rounding. Left: 6 µm z-slice showing macrophages entering the germband without a mitotically rounding cell at the entry site upon *ß-integrin RNAi* expression in the ectoderm. Right: 3-D rendering of a 26 µm z-stack of the same embryo with all mitotically rounded cells in the field of view segmented in grey and the first macrophage nucleus in yellow. Both lateral view. Macrophage nuclei *srpHemo-H2A::3xmCherry* (magenta), membranes Resille::GFP and DE-Cad::GFP (green). Corresponds to **Fig. 4e.** Scale bar: 20µm.
